# TEX15 safeguards neuronal diversity in the mouse olfactory system

**DOI:** 10.64898/2026.05.27.728302

**Authors:** Nusrath Yusuf, David H. Brann, Chuhanwen Sun, Joshua Danoff, Alina Irvine, Hetanshi Patel, Nader Boutros Ghali, Jerome Kahiapo, Jiekun Yang, Sandeep R. Datta, P. Jeremy Wang, Kevin Monahan

## Abstract

Animals detect diverse chemicals using hundreds of chemoreceptor proteins. In many organisms, sensory chemoreceptors obey a one-receptor-type-per-cell rule, which organizes information gathered by sensory neurons. A key question remains how receptor expression gets diversified so that hundreds of chemoreceptors get expressed across the population of sensory neurons. Here we show that diverse odorant receptor (OR) expression in mice requires *testis expressed gene 15* (*Tex15*). We find that *Tex15* regulates OR expression in neuronal precursor cells. In *Tex15* deficient (*Tex15-/-*) mice, OR transcript levels are increased in precursors, a handful of early-transcribed OR genes get expressed at abnormally high levels, and these ORs come to dominate the olfactory sensory neuron population. This results in a profound reduction in the diversity of OR gene choice as well as disrupted spatial patterning of the olfactory epithelium, without disrupting the one-receptor-type-per-cell rule. Tex15 encodes a transcriptional repressor, and we show that *Tex15-/-* mice have reduced heterochromatin-associated H3K9me3 on OR genes. Thus, we propose that *Tex15* directs a heterochromatin-based transcriptional repression mechanism that prevents uneven transcriptional onset from skewing OR choice. This mechanism safeguards the cellular diversity that is essential for broadly tuned chemosensation.

## Text

Animal sensory systems deploy large families of receptor proteins to detect and distinguish among stimuli present in the environment. Often, these receptors follow a one-receptor-type-per-cell rule, with cells that express the same receptor corresponding to distinct sensory neuron subtypes^1^. This structure is epitomized by the mouse olfactory system, in which each mature olfactory sensory neuron (mOSN) transcribes only one allele of one odorant receptor (OR) gene out of 1,141 protein-coding ORs^2–5^. Transcription of one OR per cell, termed “OR choice”, defines hundreds of distinct mOSN subtypes^6,7^, each with a characteristic receptor field and downstream synaptic target in the olfactory bulb^8–10^. As a result, OR choice connects odorant-to-receptor binding to the activation of specific olfactory bulb locations, spatially organizing information from hundreds of receptors for downstream processing^11^. The gene regulatory mechanisms governing OR expression have been of keen interest since the genes were first identified^12^.

OR gene choice has two key features: singularity, the expression of one OR per cell, and diversity, the expression of the full OR gene repertoire across the OSN population. Singularity is achieved through a positive feedback loop, wherein transcription of one OR allele reinforces its own expression and suppresses transcription of all other OR genes^13–15^. Diversity is less well understood, but existing evidence suggests an important role for the epigenetic regulation of OR gene chromatin as OSNs differentiate. In differentiating OSN precursors, OR genes become compacted into heterochromatin that is marked by histone H3 lysine 9 trimethylation (H3K9me3)^16,17^. Lyons et al showed that deleting the H3K9 methyltransferases Ehmt1 (Glp) and Ehmt2 (G9a) reduces OR diversity in a dosage-dependent manner, with the strongest effect in double knockouts. However, the Ehmt1/Ehmt2 double-knockout is embryonic lethal and also disrupts singular OR choice, complicating the interpretation of this phenotype. More recently, the epigenetic repressor Trim66 was shown to bind enhancer elements for OR genes. Deletion of Trim66 impacts OR transcript levels and also causes co-expression of multiple OR genes per cell. Thus, epigenetic silencing of OR transcription in precursors appears critical for diverse OR expression, but it has not been possible to separate diversification from receptor choice.

To identify regulatory proteins that enable diverse OR gene choice, we examined the expression of heterochromatin associated genes in RNA-seq data from staged OSN precursors^14^. We focused on genes that are upregulated at the transition from globose basal cells (GBCs) to immediate neuronal precursors (INPs), which corresponds to the onset of OR transcription and to the onset of H3K9me3 deposition on OR genes^17^. In total, we identified 2,070 genes that are significantly upregulated in INPs relative to GBCs, of which 14 are associated with the geno ontology term heterochromatin formation (Extended Data Fig. 1a-b). Among these, *testis expressed gene 15* (*Tex15*), exhibits the largest increase in expression, with a 35-fold increase in transcript levels at the INP stage. We next examined *Tex15* expression across differentiation. *Tex15* transcript levels are very low in olfactory stem cells (Horizontal Basal Cells and GBCs), high in INPs and iOSNs, and then low again in mOSNs (Fig. 1a). We confirmed this at the protein level by immunostaining postnatal day 7 (P7) MOE tissue sections from mice bearing an Ngn1-GFP reporter, which labels INPs and iOSNs^17^. This revealed TEX15 immunoreactivity in the nuclei of basal GFP+ cells (INPs), but not in the more apical GFP+ cells (iOSNs) or the OMP+ cells (mOSNs) (Fig. 1b, Extended Data Fig. 1c).

**Figure 1.**
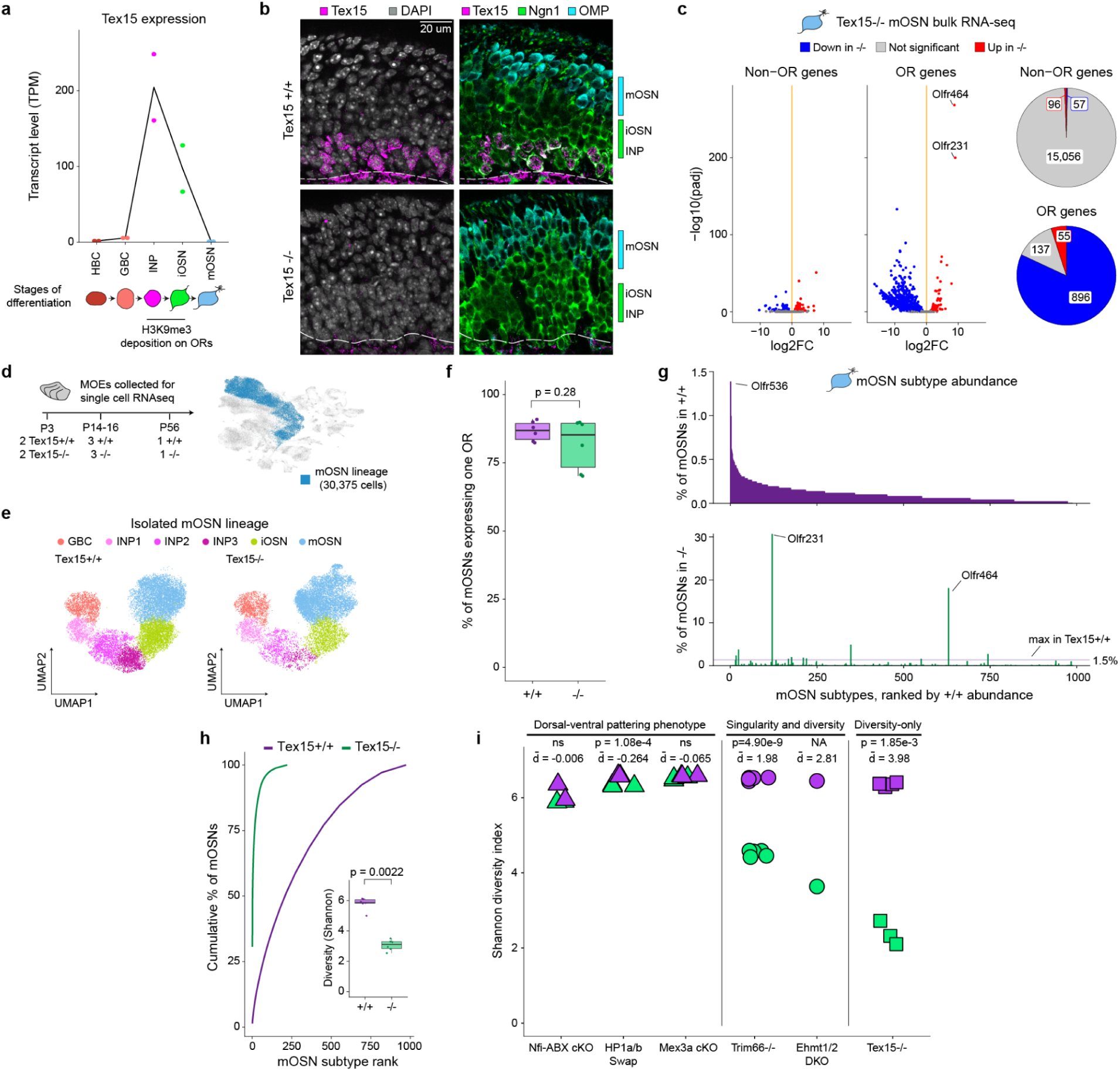
Extreme loss of mOSN diversity in Tex15-/- mice. **a**, Tex15 transcript levels, in transcripts per million (TPM), in cells purified at different stages of mOSN differentiation. **b,** Immunohistochemistry for Tex15 (magenta), Ngn1-GFP (green - INPs and iOSNs), and OMP (cyan - mOSNs), in MOE tissue sections. Nuclei are labeled with DAPI (gray). **c,** (left) Fold change in transcript levels between Tex15-/- (n=3) and Tex15+/+ mOSNs (n=5). ORs and non-olfactory receptor genes are plotted separately. (right) Summary of differentially expressed genes from each group (padj < 0.05 for a >50% change in expression, Wald test). **d,** (left) MOE tissue samples analyzed by single-cell RNA-seq. (right) UMAP projection of cells from all 12 libraries, after filtering and integration. Cells in the mOSN lineage are labelled in blue. **e,** Cells in the mOSN lineage after reclustering and assignment to cell stages. The INP population is divided into three stages, INP1-INP3. Tex15+/+ = 16,832 cells, Tex15-/- = 13,543 cells. **e,** Cells across the neuronal lineage colored by genotype - Tex15+/+ in purple, Tex15-/- in green. **f,** The percentage of mOSNs expressing a single OR, using a threshold of 3 or more unique transcripts, is not significantly different between Tex15+/+ and Tex15-/- samples (p = 0.28, t-test). **g,** Abundance of individual mOSN subtypes (mOSNs expressing the same OR), ranked by abundance in Tex15+/+ MOEs. The maximum value for wildtype mOSNs (top) is indicated by a dotted line for Tex15-/- mOSNs (bottom). **h,** Cumulative fraction of mOSNs accounted for by each mOSN subtype, ranked by abundance within each genotype. Inset, the diversity (Shannon diversity index) of the Tex15-/- mOSN population is significantly reduced compared to the Tex15+/+ (p=0.0022, t-test). **i,** Shannon diversity index calculated from OR transcript levels, measured by RNAseq, in several different knockouts and/or null studies - d̄ indicates difference in mean for control vs experimental sample.

TEX15 encodes a transcriptional repressor that has previously been shown to repress transcription of transposable elements (TEs) in the male germline. Mice with a Tex15 mutant allele (Tex15-/-), in which most of the coding sequence is replaced by a beta galactosidase-neomycin cassette, exhibit increased TE expression in male germ cells, causing arrested spermatogenesis and infertility^18^. A similar phenotype is observed in mice in which just the N-terminal “DUF3715” domain of Tex15 has been deleted^19^. Notably, DUF3715 is also found in TASOR1 and TASOR2, which are core components of HUSH (Human Silencing Hub) complexes that represses TE transcription in somatic cells. Like Tex15, TASOR1/TASOR2 require DUF3715 for their repressive activity, and domain-swap experiments show that TEX15’s DUF3715 domain can also suffice^20^. Consistent with its role in mice, mutations in *Tex15* have also been linked to male infertility in humans^21–23^. However, the function of TEX15 outside the male germ line has not been explored.

To investigate the function of TEX15 in the olfactory system, we examined Tex15-/- mice. As a first step, we evaluated the overall structure and cellular composition of the main olfactory epithelium (MOE) in *Tex15-/-* mice bearing Neurog1-GFP. Tissue sections from *Tex15-/-* mice lack TEX15 immunoreactivity in INPs, leaving only non-specific staining in more basal mesenchymal cells (Fig. 1b). Otherwise, the overall appearance of MOE sections from *Tex15-/-* mice is normal, with distinct layers of Neurog1-GFP+ and OMP+ cells.

### Loss of mOSN diversity in *Tex15-/-* mice

Next we asked whether the absence of Tex15 affects OR gene expression by mOSNs. To accomplish this, we crossed the *Tex15-/-* allele together with an Omp-GFP reporter allele^24^, which labels mOSNs, and purified mOSNs by FACS for RNA-seq. Comparison of *Tex15-/-* mOSNs to data from *Tex15* wildtype mOSNs^25^ revealed widespread changes in OR transcript levels but limited changes in non-OR gene expression (Fig. 1c). The vast majority of protein-coding OR genes are differentially expressed in *Tex15-/-* mOSNs, with 55 (5.06%) OR genes significantly upregulated and 896 (82.35%) significantly downregulated. Many of the downregulated ORs are lost completely; 786 are no longer detected in *Tex15-/-* mOSNs (<0.1 transcripts per million/TPM). OR genes fall into two families, the clustered Class I and the larger, dispersed Class II subfamily; both show a similar effect (Extended Data Fig. 1d). Strikingly, a pair of class II OR genes, *Olfr231* and *Olfr464*, are upregulated more than 250-fold. These transcript levels far exceed that of any OR gene in wildtype mOSNs (Extended Data Fig. 1e). Notably, the same two ORs are also among the top expressed ORs in Ehmt1/Ehmt2 double-knockout mice^26^, suggesting a link between TEX15 and H3K9 methylation. We also observe downregulation of the trace amine associate receptors (TAARs), another chemoreceptor family expressed by mOSNs, (Extended Data Fig. 1f)^27^. However, unlike in the male germline, TE transcript levels remain largely unchanged, and the few significant changes observed are bidirectional rather than a broad upregulation (Extended Data Fig 1g).

We next examined whether these alterations in OR transcript levels correspond to changed abundance of mOSN subtypes. To address this, we used single-cell RNA-seq (scRNAseq) to analyze the MOEs at multiple ages – postnatal day 3 (P3), postnatal day 14-16 (P14-16), and adult (P56), capturing cells spanning mOSN differentiation (Fig. 1d). We integrated data from 6 pairs of *Tex15-/-* mice and wildtype littermate controls and then isolated and re-clustered the cells in the OSN lineage (*Tex15+/+*= 16,832 cells, *Tex15-/-*=13,543 cells). Expression of cell stage-specific marker genes is similar in both genotypes (Extended Data Fig. 2a), allowing us to define developmental stages using the combined data set, including three substages within the INP population (INP1-3) (Fig. 1e). In our data, *Tex15* transcript levels peak in INP3s, which are greatly reduced in *Tex15-/-* cells, with the remaining transcripts presumably reflecting transcription through the beta galactosidase cassette (Extended Data Fig. 2b). Notably, Tex15-/- MOEs show a shift toward differentiation, with significantly fewer immature cells (INP3 and iOSNs) and more mOSNs (Fig. 1e, Extended Data Fig. 2c). Since OR choice regulates OSN maturation^15,26^, this raises the possibility that cells may choose an OR gene more quickly in *Tex15-/-* mice.

We next examined OR expression in *Tex15-/-* mOSNs. Critically, the vast majority of *Tex15-/-* mOSNs still express only one OR per cell, and OR transcript levels per cell are similar to *Tex15+/+* mOSNs (Fig. 1f, Extended Data Fig. 2d). After excluding mOSNs expressing multiple ORs, we assigned each cell a subtype based on its expressed OR. In *Tex15+/+* mice, we see a broad distribution of mOSN subtypes with the most abundant subtype, *Olfr536* mOSNs, accounting for 1.38% of the total (Fig. 1g). In contrast, *Olfr231* and *Olfr464* together account for nearly half of the mOSNs in *Tex15-/-* mice: 30.7% and 18.0%, respectively. Overall, the *Tex15-/-* results in an mOSN population with fewer subtypes but more cells per subtype, leading to a dramatic reduction in the diversity of the mOSN repertoire (Fig. 1h).

The loss of diversity observed in Tex15-/- mice is distinct from other mutations that affect OR gene choice (Fig.1i). We compared OR transcriptomes diversity across several mutant RNAseq data sets using the Shannon Diversity Index. Mutations that affect heterochromatin deposition (Nfia-abx deletion) or function (Cbx-swap) have previously been shown to alter OR dorsal ventral patterning, but these have little or no effect on OR transcriptome diversity. Deletion of an RNA-binding protein, Mex3a, was also reported to regulate OR translation and the progression to OR choice, and its loss strongly affects ventral OR expression, but loss of Mex3a does not significantly impact overall OR diversity. In contrast, mutations that result in OR co-expression, including deletion of transcriptional repressor Trim66 and combined deletion of Ehhmt1 and Ehmt2 also exhibit reduced transcriptome diversity. Only loss of Tex15 reduces diversity without affecting singularity, and the effect of Tex15-/- on OR diversity is much more pronounced than either Trim66 or Ehmt1/2. Thus, to our knowledge, loss of Tex15 is the only mutation that strongly affects OSN diversity without also leading to loss of OR singularity.

### Early-expressed ORs dominate in *Tex15-/-* mice

How does *Tex15* diversify OR choice? Based on its expression timing, we hypothesized that Tex15 regulates OR transcription in precursors. To test this, we used our scRNA-seq data to examine OR expression at each stage of differentiation. Tex15 expression peaks in INP3s (Fig. 2a, Extended Data Fig. 2b), which aligns with the peak of OR co-expression prior to choice, using a threshold of 1 transcript per cell (Fig. 2b). In *Tex15-/-* mOSNs, the rate of OR co-expression is significantly increased, with more OR genes detected per cell (Fig. 2b, Extended Data Fig. 3a). Moreover, we observe that most *Tex15-/-* INP3s express multiple OR genes using a higher threshold of 3 or more transcripts per cell, whereas wildtype INP3s rarely express OR genes at this level (Fig. 2c, Extended Data Fig. 3b). Finally, we observe a significant increase in the total number of OR transcripts detected per cell in *Tex15-/-* INP3s (Fig. 2d, Extended Data Fig. 3c). All of these metrics are similar between *Tex15-/-* and *+/+* at the other cell stages (p > 0.05, post hoc t-test). Taken together, these findings demonstrate that the loss of *Tex15* leads to increased OR gene transcription specifically in INP3s.

**Figure 2:**
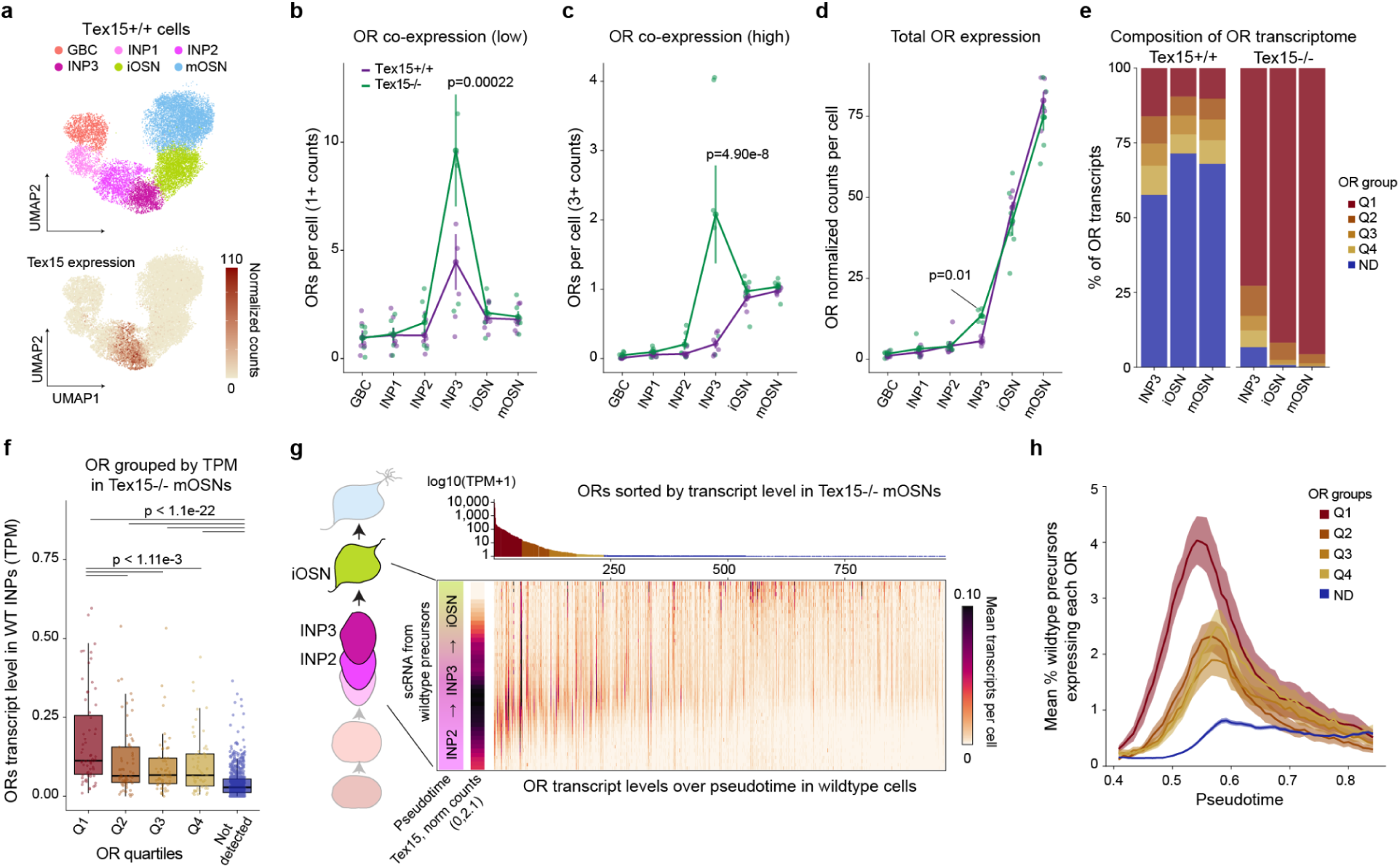
Early-expressed ORs dominate in Tex15-/- mice. **a**, Tex15 transcripts peak in INP2/INP3 cells in Tex15+/+ mice. **b,** Tex15-/- INP3s express more OR genes per cell using a low threshold (1 or more OR counts), which detects increased low-level co-expression (p=0.00022, post-hoc t-test, mixed linear model ANOVA). **c,** On average, Tex15-/- INP3s express multiple OR genes at a high level (3 or more counts), which is rarely observed in Tex15+/+ INP3s (p=4.90e-8, post-hoc t-test, mixed linear model ANOVA). **d,** Normalized OR counts per cell, which detects atypical high OR expression in Tex15-/- INP3s (p=0.01, post-hoc t-test, mixed linear model ANOVA). **e,** Relative abundance of OR transcripts, grouped according to the OR expression groups. ORs are split into five groups: ORs expressed in Tex15-/- are split into four quartiles (n=75 to 76 each, total 302) by expression in Tex15-/- and a “not detected” OR group (ND), which are present at less than 0.1 TPM (n=786). **f,** In wildtype INP bulk RNA-seq, the Q1 ORs exhibit significantly higher median transcript levels than the Q2 (p=0.0298), Q3 (p=0.00132), Q4, and ND ORs (adjusted p-values, Dunn test with correction for multiple testing). **g,** (left) scRNA-seq data from 154,642 wildtype OSN precursors from the INP2 to iOSN stages (Brann et al 2026) was binned based on the pseudotime score for each cell. (right) Mean transcript levels for Tex15 and each OR over binned pseudotime. ORs are arranged by their level of expression in Tex15-/- mOSNs (top). **h,** For each expression group, the mean percentage of cells detected expressing each OR over pseudotime. Line shows mean, shaded area shows SEM. For a-c, points show mean and SEM across Tex15+/+ and Tex15-/- samples (n=6 for each). P-values were calculated from post-hoc t-tests on an ANOVA comparing genotypes for each cell population.

Next, we asked how the altered OR transcript levels we observe in *Tex15-/-* precursors are related to subsequent OR choice. We split the OR genes into quartiles based upon their transcript levels in *Tex15-/-* mOSNs (Q1-4, n= 75 to 76 ORs for each), and defined an additional set containing all ORs that are not detected (ND) (TPM < 0.1, n=786 ORs) (Extended Data Fig. 3d). In our scRNAseq data, the composition of OR transcripts in *Tex15-/-* INP3s is biased towards the *Tex15-*expressed ORs, and this bias becomes more prominent as differentiation continues to the iOSN and mOSN stages (Fig. 2e). However, OR transcript levels are low in these cells, limiting our ability to quantify individual OR expression. To provide a more sensitive test of this relationship, we examined gene expression in FACS purified Ngn1-GFP+ cells from *Tex15-/-* and *Tex15* heterozygous mice. For these experiments, we collected GFP+ cells from adult mice 10-days after using methimazole lesioning to induce MOE regeneration. RNAseq revealed pervasive changes in OR gene expression in *Tex15-/-* mice, with ORs accounting for most of the differentially expressed genes (Extended Data Fig. 3e-f). Critically, OR expression changes in Ngn1-GFP+ cells are highly correlated with those in OMP-GFP+ mOSNs, consistent with altered precursor OR expression skewing the resulting mOSN population (Extended Data Fig. 3g).

Why does altered OR expression in *Tex15-/-* precursors favor specific OR genes? We hypothesized that some OR genes have a transcriptional advantage in INPs that is normally suppressed but revealed in *Tex15-/-*. To test this hypothesis, we examined how OR expression in *Tex15-/-* mOSNs relates to OR expression in *Tex15+/+* INPs^14^. Strikingly, the top quartile of OR genes in *Tex15-/-* mOSNs (Q1) are expressed at significantly higher levels in wildtype INPs than the other groups, while ORs not detected in the *Tex15-/-* (ND) are expressed at significantly lower levels (Fig. 2f). Thus, OR choice in *Tex15-/-* mice reflects OR transcription in wildtype INPs.

Next we examined whether choice in *Tex15-/-* mice relates to the timing of OR transcription within the INP stage. We analyzed data from a large scRNA-seq dataset from wildtype OSN differentiation, which includes 154,642 cells from the interval spanning the onset of OR transcription to OR choice^28^. Using pseudotime values, we grouped cells from the Class II OR lineage into bins and calculated the mean transcript level of each OR per bin. This analysis revealed a striking relationship between OR transcript levels in *Tex15-/-* mOSNs and OR transcription in precursors (Fig. 2g). ORs that are highly expressed by *Tex15-/-* mOSNs exhibit a wave of strong transcription in early precursors, which subsides prior to choice. Notably, the peak of this wave aligns with the peak of *Tex15* transcript levels. Next, we examined the breadth of OR expression across this population of cells. We calculated the percentage of cells expressing each OR per bin, examining how this varied by group over pseudotime. ORs from Q1 are expressed by a higher percentage of cells, and this expression occurs earlier in pseudotime than ORs from the other groups (Fig. 2h). The prevalence of cells expressing Q1-Q4 ORs decreases in later bins, indicating that transcription is normally repressed after the initial pulse. As a result of this decrease, the expression rates for OR genes from all 5 groups ultimately reach similar levels in iOSNs. Accordingly, we calculated each OR’s mean pseudotime value, summarizing its transcription timing before singular choice, and observed the same relationship between *Tex15-/-* transcript levels and early expression (Extended Data Fig. 3h). Therefore, TEX15 loss favors the choice of early-expressed OR genes that are normally repressed in wildtype precursors.

### Assembly of enhancer hubs around the dominant OR genes

Wildtype precursors express *Olfr231* very early, even when compared to other Q1 ORs, which may contribute to its dominance in *Tex15-/-* mice (Extended Data Fig. 3i). In contrast, the other dominant OR in *Tex15-/-* MOEs, *Olfr464*, resembles other Q1 ORs. However, *Olfr464* is distinctive in a different way: it is located between a pair of Greek Islands (GIs) (Extended Data Fig. 3j), which are specialized enhancer elements that control the choice of one Class II OR gene by assembling into an interchromosomal (*trans*) enhancer hub around the chosen OR gene (Fig. 3a)^25,29^. Indeed, GI proximity is associated with Class II OR expression in *Tex15-/-*, with the top 3 quartiles significantly closer to GIs than ND ORs (Fig. 3b). Thus, we hypothesized that the dominant OR genes, *Olfr231* and *Olfr464*, preferentially assemble into GI enhancer hubs, stabilizing their choice. To test this, we FACS purified *Tex15-/-* mOSNs and performed in situ Hi-C, comparing the results to existing Hi-C data from *Tex15* wildtype mOSNs^25^. We first examined whether OR genes and OR enhancers form interchromosomal hubs in *Tex15-/-* mOSNs (Fig. 3c-d). The overall pattern of interactions among OR genes and GIs is similar to wildtype, though with a small reduction in mean *trans* contact strength. Otherwise, global features of chromosome folding are preserved in *Tex15-/-* mOSNs. For example, compartment scores are highly correlated between the two conditions (Extended Data Fig. 4a), and although OR-containing regions show a small increase, most ORs remain in the silent “B” compartment (Extended Data Fig. 4b). Notably, compartment scores around upregulated ORs, such as *Olfr464*, are unchanged, consistent with the observation that they remain inactive in most cells (Extended Data Fig. 4c-e).

**Figure 3:**
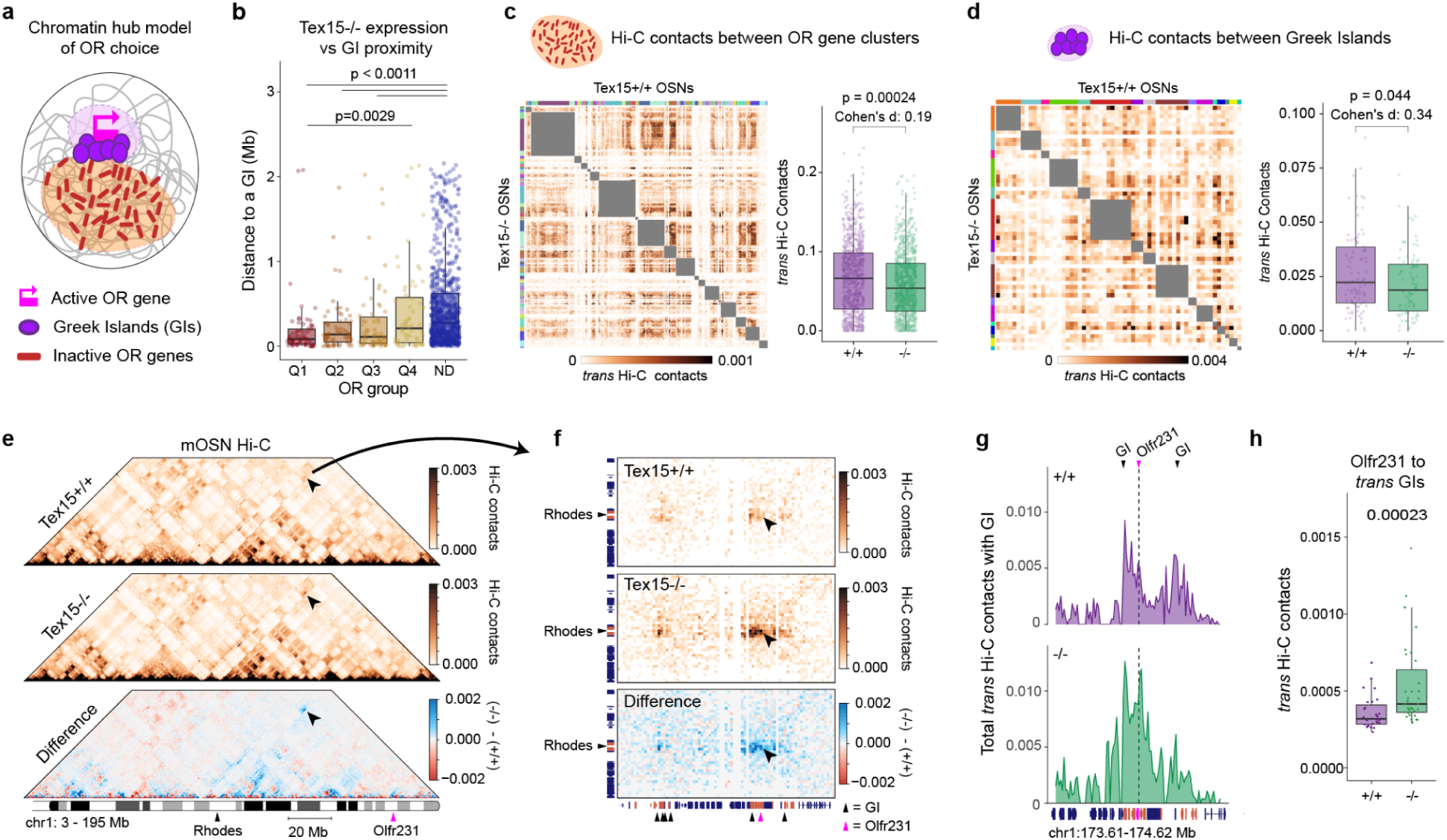
Assembly of enhancer hubs around the dominant OR genes. **a**, a Greek Island (GI) enhancer hub assembles around the active OR gene. **b,** The top 3 quartiles of OR genes in the Tex15-/- are significantly closer to GIs than the ND OR genes, and Q1 is also closer than Q4 (Dunn test, corrected for multiple comparisons). **c,** (left) Heatmap of interchromosomal (*trans*) Hi-C contacts between OR clusters, binned at 25 Kb resolution. Tex15+/+ data is above the diagonal, Tex15-/- below. Interactions between bins located on the same chromosome are masked (gray). Color bars indicate OR clusters. (right) Quantification of *trans* contacts between OR cluster bins reveal significant but marginal reduction in contacts in Tex15-/- OSNs (p = 0.00024, Wilcoxon, Cohen’s d = 0.19). **d,** (left) Heatmap of interchromosomal (*trans*) Hi-C contacts between Greek Islands (GIs),as in **c**, but color bars indicate chromosome. (right) Quantification of interchromosomal (*trans*) contacts between GIs reveals a small but significant reduction in contacts (p = 0.044, Wilcoxon, Cohen’s d = 0.34). **e,** The difference in Hi-C contacts between Tex15-/- and Tex15+/+ mOSNs across a 105 Mb region of chromosome 1. Triangles indicate the location of *Olfr231* and a distant GI, Rhodes. Arrowhead indicates contacts between these loci. 500 Kb resolution **f,** Zoom-in of Hi-C contacts between the Rhodes locus and the *Olfr231* locus, which includes multiple nearby GIs. OR genes are coral, non-OR genes are navy. Arrowheads indicate contacts between Rhodes and Olfr231 at 25 Kb resolution. **g,** Summed interchromosomal (*trans*) contacts from GIs to the *Olfr231* locus. 10 Kb resolution. **h,** *trans* Hi-C contacts between *Olfr231* and each *trans* GI (Wilcoxon, p=0.00023).Panels **c-d**, use balanced Hi-C contacts calculated after merging data from two biological replicates.

Having confirmed that *Tex15-/-* mOSNs form GI enhancer hubs, we next asked whether there is increased association of *Olfr231* and *Olfr464* with these structures. For Olfr231, we observe increased long-range Hi-C contacts with Rhodes, a GI enhancer located 81.6 Mb away (Fig. 3e). The increase in Hi-C contacts is strongest over *Olfr231* itself, but also evident on several GIs located near it (Fig. 3f). *Olfr231* also shows significantly increased Hi-C contacts with GIs on different chromosomes, indicating formation of an interchromosomal enhancer hub around it (Fig. 3g-h). For *Olfr464*, the locus already makes strong contacts with GIs in wildtype cells due to the presence of a pair of nearby GIs (Extended Data Fig. 3d). However, interchromosomal contacts to GIs increase specifically over *Olfr464* in *Tex15-/-* mOSNs (Extended Data Fig. 4e). Thus, both of the dominant ORs in *Tex15-/-* mice associate with interchromosomal GI hubs.

Even though *Olfr231* and *Olfr464* associate with GI hubs, it remains possible that an orthogonal mechanism drives their dominance in *Tex15-/-* mice. GI hubs are assembled by an adapter protein, LDB1, and deletion of *Ldb1* in mOSNs leads to loss of hubs and reduced OR transcription^25,30^. Thus we asked whether *Ldb1* is also required for OR gene expression in *Tex15-/-* mOSNs. To accomplish this, we conditionally deleted *Ldb1* using an OMP-cre allele in a *Tex15-/-* background and then profiled gene expression by RNA-seq. We then compared OR expression in double-knockout cells to OMP-cre controls and previously reported OMP-cre *Ldb1* knockouts^25^. Consistent with enhancer hub dependence, loss of Ldb1 significantly reduces aggregate OR transcript levels *Tex15/Ldb1* double KO mOSNs, as in *Ldb1*KO mOSNs (Fig. 4a-b). In contrast, the double KO has a much stronger effect on OR transcriptome diversity (Fig. 4c).

**Figure 4:**
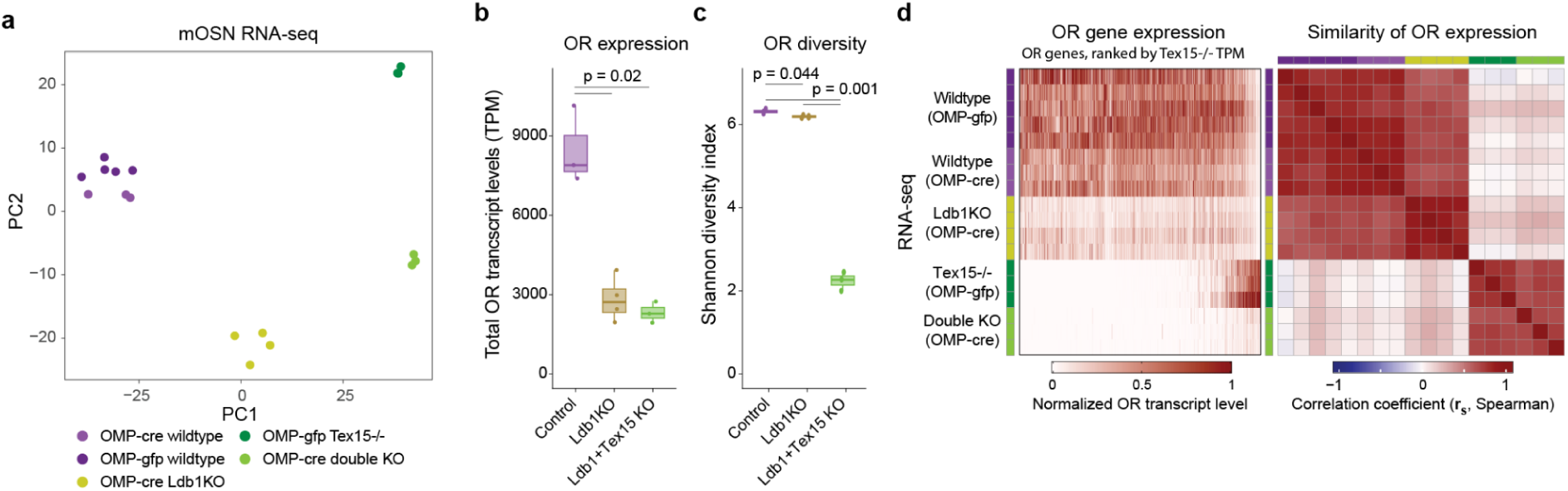
OR expression still depends on Ldb1 in Tex15-/- OSNs. **a**, Principal component analysis of bulk-RNA-seq datasets analyzed together with the Tex15 and Ldb1 double knockout. Wildtype mOSN labeled with either OMP-Cre and OMP-GFP are similar to each other, otherwise samples from each genotype cluster together as a discrete group. All Hi-C analyses (a-i) show balanced *trans* Hi-C contacts calculated after merging data from two biological replicates. **b,** Total OR transcript levels are significantly reduced in Ldb1KO OSNs (n=4, from Monahan et al 2019), and Tex15/Ldb1 double-knockout OSNs libraries (n=3) compared to control OMP-cre labeled mOSNs (n=3). (t-test, corrected for multiple comparisons) **c,** Diversity of OR transcripts is significantly reduced in double KO mOSN relative to control (p=0.001, Cohen’s d = 25.0) than the loss of Ldb1 alone relative to control (p=0.047, Cohen’s d = 2.79)(t-test, corrected for multiple comparisons). **d,** (left) OR transcript levels across mOSNs with different genotypes. Each row is a biological replicate. Transcript levels are normalized for each OR gene. ORs are sorted by their transcript levels in Tex15-/-;OMP-gfp mOSNs. (right) Spearman correlation between OR transcript levels across samples.

Strikingly, double KO mOSNs retain the skewed OR expression pattern characteristic of *Tex15-/-* cells but at the reduced overall transcript levels characteristic of *Ldb1*KO cells (Fig. 4d). This indicates that the identity of the chosen OR, determined upstream by the *Tex15*-dependent diversity mechanism, is unaffected by disrupting the hub that sustains its expression. Overall, these results indicate that the *Tex15-/-* biases OR choice without compromising the singularity mechanism, and that these two processes can be genetically dissociated.

### TEX15 directs de novo deposition of H3K9me3 on OR genes

Since reduced diversity is seen in H3K9-methyltransferase double knockouts^31^, we hypothesized that TEX15 is involved with directing heterochromatin formation on OR gene clusters. To test this, we performed native ChIP-seq for H3K9me3 in adult MOE tissue from *Tex15-/-* mice and wildtype controls (Fig. 5a). As expected, we observe reduced H3K9me3 signal on OR genes in *Tex15-/-* tissue (Fig. 5b, Extended Data Fig. 5a-b). Genomewide, H3K9me3 reductions are highly localized to OR gene clusters, standing out against the background of whole chromosomes (Fig. 5c). Differential enrichment analysis confirms that the *Tex15-/-* particularly impacts H3K9me3 on ORs, with OR loci accounting for the vast majority of significantly depleted regions (Fig. 5d). Moreover, as a group, H3K9me3 regions that overlap ORs exhibit a strong reduction in signal relative to the rest of the genome (Fig. 5e). Examination of the H3K9me3 signal on the body of each OR gene does not reveal a strong relationship between fold change in H3K9me3 and OR expression in Tex15KO OSNs (Fig. 5f). While the ND ORs do exhibit significantly greater reduction relative to the Q1 and Q3 groups, the effect sizes are small.

**Figure 5:**
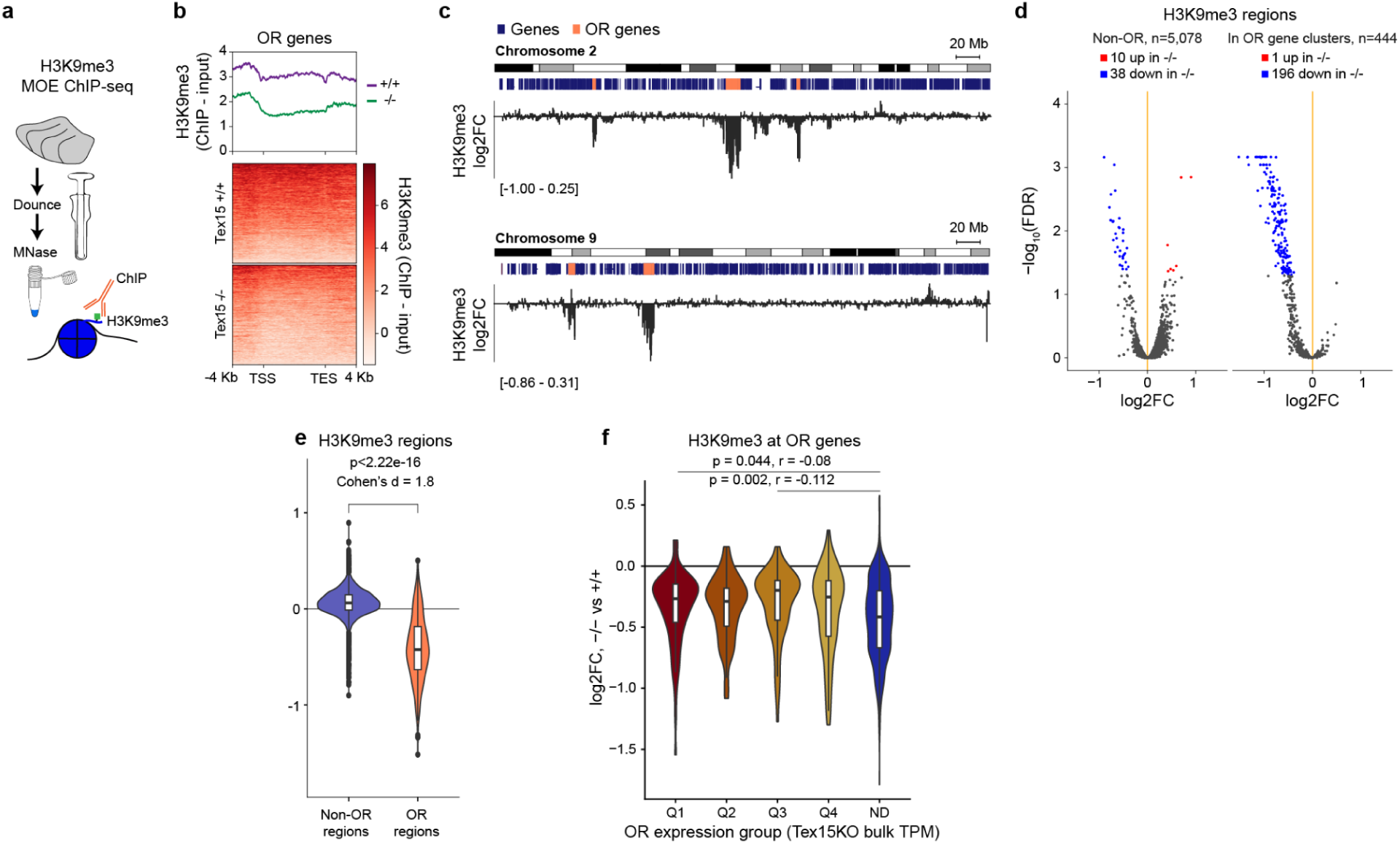
Tex15 directs de novo deposition of H3K9me3 on OR genes. **a**, H3K9me3 ChIP-seq from MOE of adult Tex15-/- mice and wildtype controls. **b,** Heatmap and plot showing H3K9me3 signal over OR genes for a representative experiment. **c,** Fold change in H3K9me3 ChIPseq signal over the entirety of chromosome 2 and chromosome 9, showing decreased signal over OR genes (coral) compared to all other genes (blue) for a representative experiment. **d,** Differential enrichment analysis of H3K9me3 regions, split based on whether they overlap OR gene clusters. Regions with a significant increase in H3K9me3 signal are in red, decrease in blue (FDR < 0.05, n=4 per condition). **d,** Fold change in ChIP signal in H3K9me3 regions, grouped based on OR cluster overlap. OR regions show a significant reduction in H3K9me3 signal compared to non-OR regions (p < 2.22e-16, t-test, Cohen’s d = 2.04). **f.** Log2fold change of H3k9me3 deposition at OR groups separated by quartiles according to their expression in Tex15-/-. ND OR have a significantly stronger reduction than Q1 and Q3 ORs (Kruskal-Walli test, post hoc Dunn Test).

Since OR genes normally gain H3K9me3 over differentiation, we hypothesized that the reductions we observe in *Tex15-/-* MOE result from impaired H3K9me3 deposition. To test this, we reanalyzed published H3K9me3 data from INPs and mOSNs (Extended Data Fig. 5c-e)^17^, and then compared changes over differentiation to changes in *Tex15-/-* mice. Consistent with our hypothesis, reductions in H3K9me3 in the *Tex15-/-* MOE are almost entirely restricted to regions that gain H3K9me3 over differentiation (Extended Data Fig. 5f). Moreover, regions that gain H3K9me3 over differentiation and overlap ORs show reduced signal in *Tex15-/-*, unlike corresponding non-OR regions (Extended Data Fig. 5g). Taken together, these findings suggest that *Tex15-/-* impairs H3K9me3 deposition over OR genes during differentiation.

### *Tex15-/-* disrupts OR spatial patterning

H3K9me3 has also been linked to the spatial patterning of the MOE. Each mOSN subtype is most abundant at a specific spatial position along the MOE’s dorsal-ventral (DV) axis (Fig 6a)^6,8,28,32,33^. Intriguingly, subtype DV position is linked to the level of H3K9me3 at the corresponding OR locus — dorsal ORs carry higher H3K9me3 than ventral ORs — and mutations that perturb heterochromatin shift the relative expression of dorsal and ventral ORs^17,28,34^.

**Figure 6:**
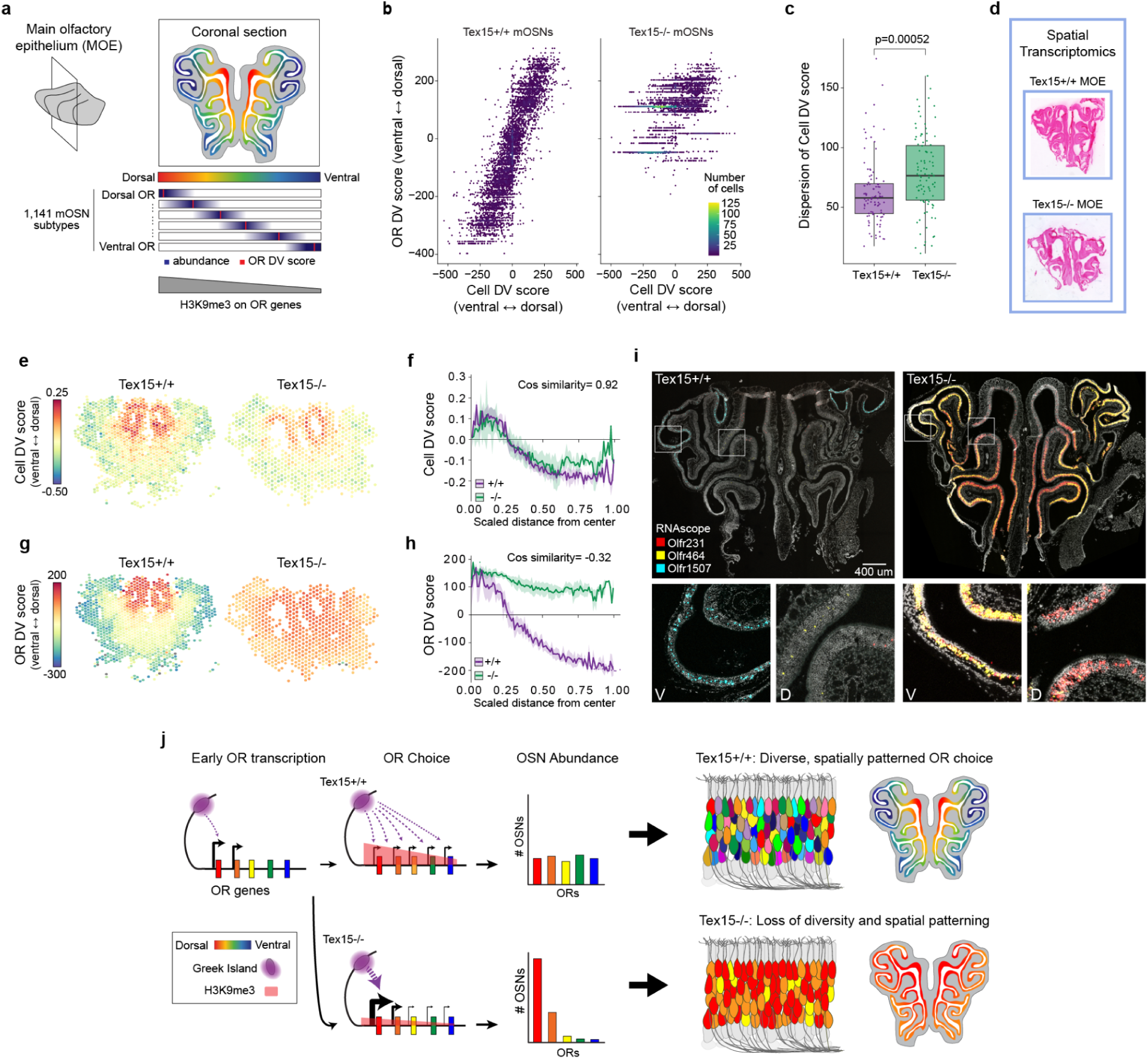
Tex15-/- results in disrupted OR spatial patterning. **a**, mOSNs subtypes are most abundant (navy) at specific positions along the dorsal-ventral (DV) axis of the MOE, summarized as an OR-DV score (red). DV score correlates with H3K9me3 levels on OR genes. **b,** The density of - mOSNs observed across the range of Cell DV scores (based on non-OR genes) and OR DV scores. **c,** Tex15-/- mOSN subtypes exhibit an increased dispersion of Cell DV scores observed within mOSN subtypes (only subtypes with 3+ cells in both conditions, p=0.00052, t-test). **d,** 10x Visium spatial transcriptomics analysis of HE-stained coronal sections of Tex15+/+ (n=6) and Tex15-/- (n=3) MOEs. **e,** Cell DV score across a representative Tex15+/+ and Tex15-/- tissue section. Cell DV score is calculated from transcript levels of dorsally and ventrally patterned genes within each 55 µm spot. **f,** Cell DV scores versus radial position. Line indicates mean across tissue sections and shaded area indicates SEM. Cos similarity measures how similar the curves are. **g,** OR DV score across a representative Tex15+/+ and Tex15-/- tissue section. OR DV Score is calculated from OR transcript levels within each 55 µm spot. **h,** OR DV scores versus radial position. Line indicates mean across slides and shaded area indicates SEM. **i,** (top) RNA Scope detection of *Olfr231* (red), *Olfr464* (yellow), and *Olfr1507* (cyan) transcripts in MOE tissue sections. Bar = 400 µm. (bottom) Larger images of boxed regions from above. **j,** Model for how Tex15 safeguards mOSN diversity by repressing the transcription of early expressed OR genes. Transcriptional repression and heterochromatin formation equalize OR transcript levels in late precursors, allowing for diverse OR choice. In Tex15-/- mice, heterochromatin formation is less efficient and OR transcription in INPs increases rapidly, thereby skewing enhancer hub assembly and choice towards the early expressed ORs.

Since the *Tex15-/-* reduces H3K9me3 deposition, we asked whether it also impacted the DV patterning of the MOE. First, we used our scRNAseq data to examine the relationship between OR choice and spatially patterned non-OR gene expression. For each mOSN, we assigned an “OR DV score” based on the typical DV position of the expressed OR, as determined in wild-type mice by Brann et al^28^. We then compared this to a “cell DV score”, calculated using the transcript levels of non-OR genes that exhibit dorsal or ventral expression, as in Tsukahara et al^6^. In our wildtype mOSNs, we observe the expected correlation between the cell DV score and OR DV score, but this relationship is disrupted in *Tex15-/-* mOSNs (Fig. 6b). In these cells, we observe a similar distribution of cell DV scores, but very few cells that express ventral ORs (Extended Data Fig. 6a,b). However, spatial patterning is not entirely lost; a correlation remains between cell DV scores for matched OSN subtypes in *Tex15-/-* and *Tex15+/+* cells (R=0.54, p=1.2e-8) (Extended Data Fig. 6c). However, dispersion of cell DV scores within each subtype is significantly increased in *Tex15-/-* (Fig. 6c). Taken together, these results indicate the coupling between spatial position and OR gene choice is weakened in *Tex15-/-* mice.

To directly examine how the *Tex15-/-* affects the spatial patterning, we used spatial transcriptomics to simultaneously map DV patterning and OR gene expression in tissue sections from adult MOEs (Fig. 6d). Examining spatially patterned non-OR genes, either individually or via a scaled cell DV score^6^, we found that the overall DV patterning of the MOE is similar between control and *Tex15-/-* mice (Fig. 6e-f, Extended Data Fig. 6d-e). However, we observe loss of ventral ORs in *Tex15-/-* mice, assessed by calculating an OR-DV score weighted by transcript abundance (Fig. 6g-h, Extended Data Fig. 6f). Consistent with our scRNAseq analysis, we observe that ventral regions of the MOE instead express dorsal ORs. This is not related to spatial variation in *Tex15* expression, as we detect TEX15 protein throughout the MOE (Extended Data Fig. 6g).

If the ventral OR genes are no longer expressed, what ORs take their place? Examination of transcript levels for individual OR genes reveals expression of *Olfr231* and *Olfr464* throughout the ventral regions of MOE sections from *Tex15-/-* mice (Extended Data Fig. 6h). To examine expression of these ORs at cellular resolution, we generated RNAscope probes for *Olfr231* and *Olfr464,* as well as *Olfr1507*, which is one of the most abundant ventral OR genes in *Tex15+/+* mice (Fig. 6i). As expected from their OR-DV scores, we observe *Olfr231* and *Olfr464* in dorsal regions of the MOE and *Olfr1507* in ventral regions in wildtype tissue sections. In contrast, *Olfr1507* is not detected in *Tex15-/-* tissue, while *Olfr231* and *Olfr464* are abundant throughout the entire ventral MOE. Thus absence of Tex15 is particularly disruptive to the choice of ventral ORs, which are broadly displaced by *Olfr231* and *Olfr464*.

## Discussion

Our findings establish that *Tex15* safeguards the diversity and spatial patterning of OR choice by repressing an early wave of OR gene transcription that occurs in mOSN precursors. During normal mOSN differentiation, a subset of ORs are transcribed earlier than others (Fig. 6j). *Tex15* is required to repress this early transcription. This repression allows the broader OR repertoire to get transcribed and places all ORs at a similar transcriptional level (Fig 2g). In *Tex15-/-* mice, repression fails and the early ORs are transcribed at abnormally high levels in INP3s. We propose that this elevated transcription allows early-expressed ORs to hijack the singular choice mechanism by driving enhancer hub assembly, as has been observed for artificially activated OR genes^14^. Since dorsal and ventral ORs exhibit systemic differences in transcriptional timing^28^, the *Tex15-/-* disrupts DV patterning, allowing a few early-expressed dorsal ORs to dominate.

We show that dysregulation of OR expression is accompanied by reduced deposition of H3K9me3 heterochromatin marks on OR loci. These marks are normally applied during differentiation, coincident with OR co-expression and choice^17^. Knocking out a pair of H3K9-directed methyltransferases (Ehmt1/2 double KO) reduces OR transcriptome diversity, similar to the loss of Tex15 but to a lesser degree (Fig. 1i). However, while *Tex15-/-* mOSNs preserve singular choice, H3K9 methyltransferase mutants exhibit co-expression of multiple ORs per cell^26^. Interestingly, mutation of *Trim66*, a transcriptional repressor that binds to Greek Islands, also results in reduced diversity and OR co-expression, similar to H3K9 methyltransferase mutants. Taken together, these findings imply that diversification and singularity are achieved through distinct mechanisms that both involve remodeling OR gene chromatin structure.

How does Tex15 regulate OR expression and heterochromatin formation on OR genes? The most parsimonious explanation for this is that these two processes are coupled by recruitment of Tex15 to OR chromatin. Tex15 lacks a predicted DNA binding domain, thus it would presumably be recruited by other factors. The proteins that TEX15 interacts with in the olfactory epithelium have not been determined, but the predicted features of TEX15 suggest a couple of possibilities. TEX15 is a large protein of 3,049 amino acids and it is dominated by predicted intrinsically disordered regions (IDRs) except for an N-terminal DUF3715 domain^35^. Notably, DUF3715 domains are only present in two other proteins in mice, TASOR and TASOR2, which are members of the HUSH1 and HUSH2 complexes, respectively. HUSH complexes silence transcription of transposable elements in somatic cells, but have also been shown to repress transcription of large gene families, such as ZNFs and clustered protocadherins (Lehner 2025; Douse et al. 2020). If TEX15 participates in an HUSH-like complex, it could be recruited to ORs by HUSH’s MPP8 subunit, which binds existing H3K9 methylated chromatin. Notably, this mechanism could explain the partial reduction in H3K9me3 at OR loci observed in *Tex15-/-* mice (Fig. 4b-d), since HUSH recruits SETDB1 which catalyzes spreading of H3K9me3 marks to nearby nucleosomes^36^. This difference between initial H3K9 methylation (by Ehmt1/2) and *Tex15*-direct spreading could also account for the differing effects of each mutant on OR singularity. A second possibility is raised by recent reports that IDRs can interact with RNA^37^. It may be that the long IDRs of Tex15 bind and sequester nascent OR transcripts, which would provide a mechanism for both recognition and repression. In any case, given Tex15’s tightly restricted expression in the MOE (Extended Data Fig. 5g), in vitro biochemical assays may be necessary to characterize this activity.

It is also possible that Tex15 regulates ORs post-transcriptionally. Transcriptomics suggests that cytosolic OR transcripts are actively degraded in INP cells^38^. This degradation has been linked to *Mex3a*, a cytosolic dual-function RNA-binding protein and E3 ubiquitin ligase. In *Mex3a* conditional knockout mice, INPs and iOSNs exhibit elevated OR transcript levels and OR co-expression, as we observe in *Tex15-/-* mice. However, loss of *Mex3a* does not affect the ratio of mOSN precursor populations, has a relatively small effect on OR transcript levels in mOSNs, and does not significantly change the overall diversity of the OR transcriptome (Fig. 1i). While quite distinct from the phenotype observed in *Tex15-/-* mice, it remains possible that *Tex15* acts on OR transcripts within the nucleus, and only indirectly affects H3K9me3 levels on OR genes.

Regardless of the specific mechanisms at work, our findings illuminate a fundamental vulnerability in the prevailing model that OR transcription initiates and drives stable OR choice through self-reinforcing positive feedback loops^13–15,17,26,39^. Recent studies have focused on characterizing the positive feedback loop that ensures that each OSN expresses only one allele of one OR gene. While positive feedback effectively ensures OR singularity, it is inherently unstable and prone to amplifying intrinsic bias. A few ORs with a transcriptional advantage can dominate the choice process, as we observe in *Tex15-/-* mice. This advantage likely comes from multiple sources. For example, some OR genes get a transcriptional head start, perhaps by lying close to a GI or having a promoter particularly active in INP cells. Indeed, we suggest that random mutations occurring over 1,000 OR promoters are likely to evolve highly active promoter sequences by chance, which may explain the early transcription of *Olfr231*. Alternatively, GI proximity may let some OR genes assemble enhancer hubs more efficiently, perhaps accounting for the frequent choice of *Olfr464*s in *Tex15-/-* mice.

Irrespective of why some ORs turn on earlier than others, our data suggests that TEX15 acts as a brake or buffer, preventing early transcription from dominating the OR choice. Consistent with this model, the presence of fewer INPs and iOSNS in *Tex15-/-* mice could result from OR choice happening faster. Accelerated choice may also account for the disruption of OR DV patterning that we observe in *Tex15-/-* mice. Ventral precursors transiently express dorsal ORs before restricting expression to ventral ORs^17,28^. The absence of a *Tex15* imposed check on OR expression may allow these dorsal ORs, particularly *Olfr231* and *Olfr464*, to dominate the ventral MOE. Finally, we note that the precipitous decline in *Tex15* expression between the INP3 and iOSN stages coincides with the switch from a transcriptional competition and spatial patterning regime to a choice regime in which positive feedback drives singular OR expression. It may be that the disappearance of TEX15 enables the progression to OR choice.

We have also recently identified a role for TEX15 in regulating vomeronasal receptor (VR) expression in vomeronasal sensory neurons (VSNs)^40^. In each system where *Tex15* function has been characterized — male germ cell TE silencing, VR expression in VSNs, and now OR expression in OSNs — it regulates large, genomically dispersed gene families. A striking feature of *Tex15* that cuts across all of these contexts is its highly restricted expression pattern. Rather than functioning as a broadly expressed transcriptional repressor, *Tex15* appears to be deployed selectively in specific cell types at specific developmental moments: spermatocytes undergoing meiosis, VSN precursors in the vomeronasal organ undergoing VR choice, and INP3 precursors in the MOE undergoing OR choice. Given its recurring role in regulating large, dispersed gene families, *Tex15* is an attractive candidate regulator in other systems where combinatorial gene expression underlies cellular identity.

---

**Extended Data 1.**
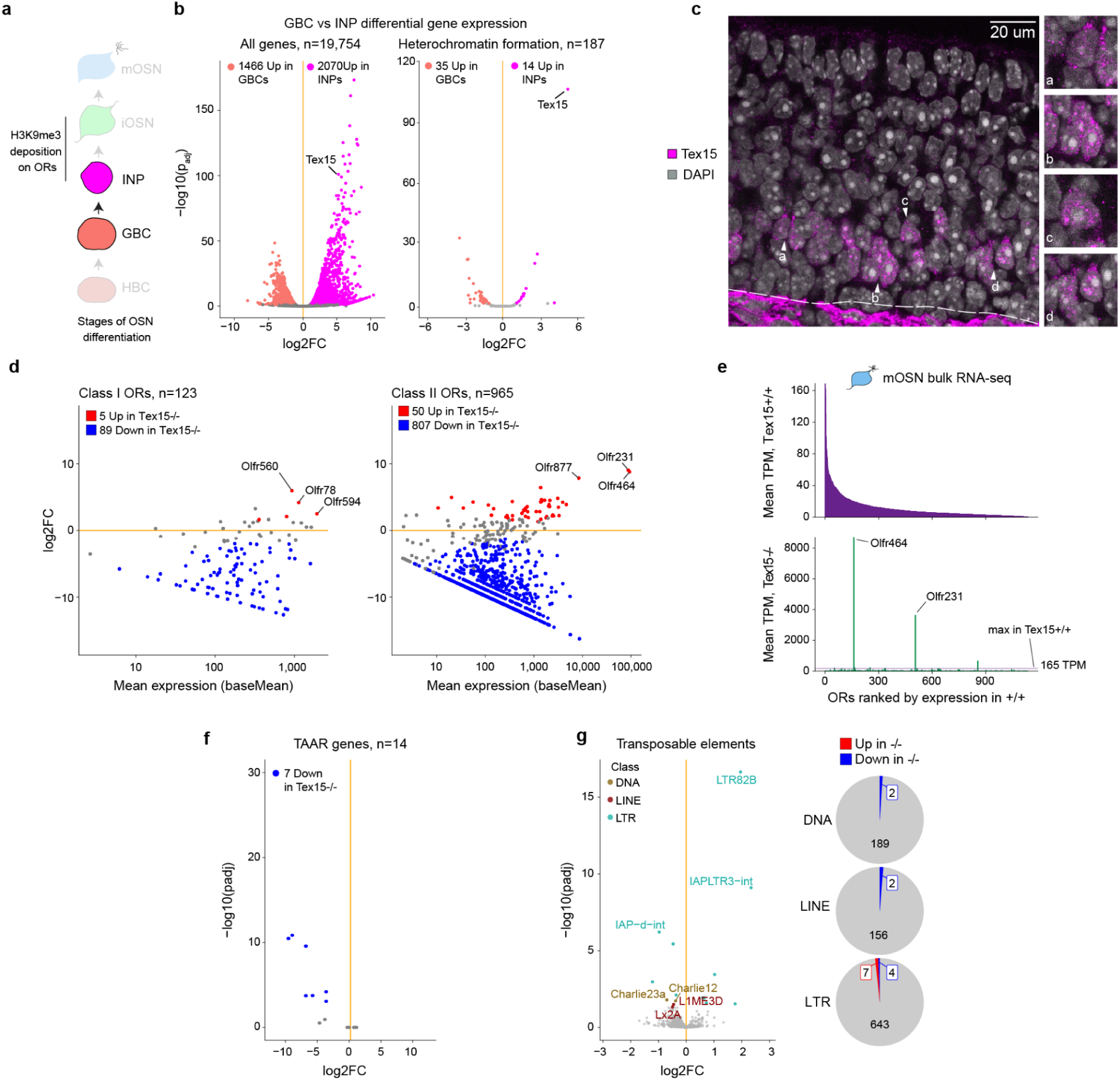
Altered receptor gene expression in the olfactory epithelium of Tex15-/- mice. **a**, Schematic illustrating the selection of stage-specific RNA-seq datasets (Pourmorady et al) used to identify factors associated with the onset of H3K9me3 deposition on OR genes. Stages: horizontal basal cells (HBC), globose basal cells (GBC), immediate neuronal precursor (INP), immature olfactory sensory neurons (iOSNs), mature olfactory sensory neurons (mOSNs). **b,** (left) Volcano plot showing differential gene expression between GBCs and INPs (n=2 per stage). Tex15 is among the genes with the largest fold change and smallest adjusted p-value. In total, 1,466 genes have significantly higher expression in GBCs and 2,070 in INPs. (right) Genes associated with the GO term “Heterochromatin formation” (n=187). 35 heterochromatin formation genes have significantly higher expression in GBCs and 14 in INPs. Tex15 is the most significantly differentially expressed gene in this set. **c,** Immunohistochemistry for Tex15 (magenta), Nuclei labeled with DAPI (gray), in MOE tissue sections with zoom-ins on cells a-d showing Tex15 protein in the nucleus. **d,** MA plots showing differential expression of Class I and Class II OR genes in Tex15+/+ (n=5) and Tex15-/- mOSNs (n=3) by RNA-seq. The top 3 upregulated ORs are labeled. **e,** Mean OR transcripts levels in Tex15+/+ and Tex15-/- mOSNs, expressed as transcripts per million (TPM). ORs are ranked by expression in Tex15+/+ mOSNs. The maximum transcript level for wildtype mOSNs (top) is indicated by a dotted line for Tex15-/- mOSNs (bottom). **f,** Volcano plot showing differential gene expression of Trace amine associated receptors (TAARs) between Tex15-/- and Tex15+/+ mOSNs. **g,** (left) Change in transposable element transcript levels between Tex15-/- and Tex15+/+ mOSNs. Significantly differentially expressed elements are color coded by class, with the top 3 from each class labeled. (right) Summary of differentially expressed TEs by repeat class. For all panels, differentially expressed genes have an adjusted p-value < 0.05 for a >50% change in expression (Wald test).

**Extended Data 2.**
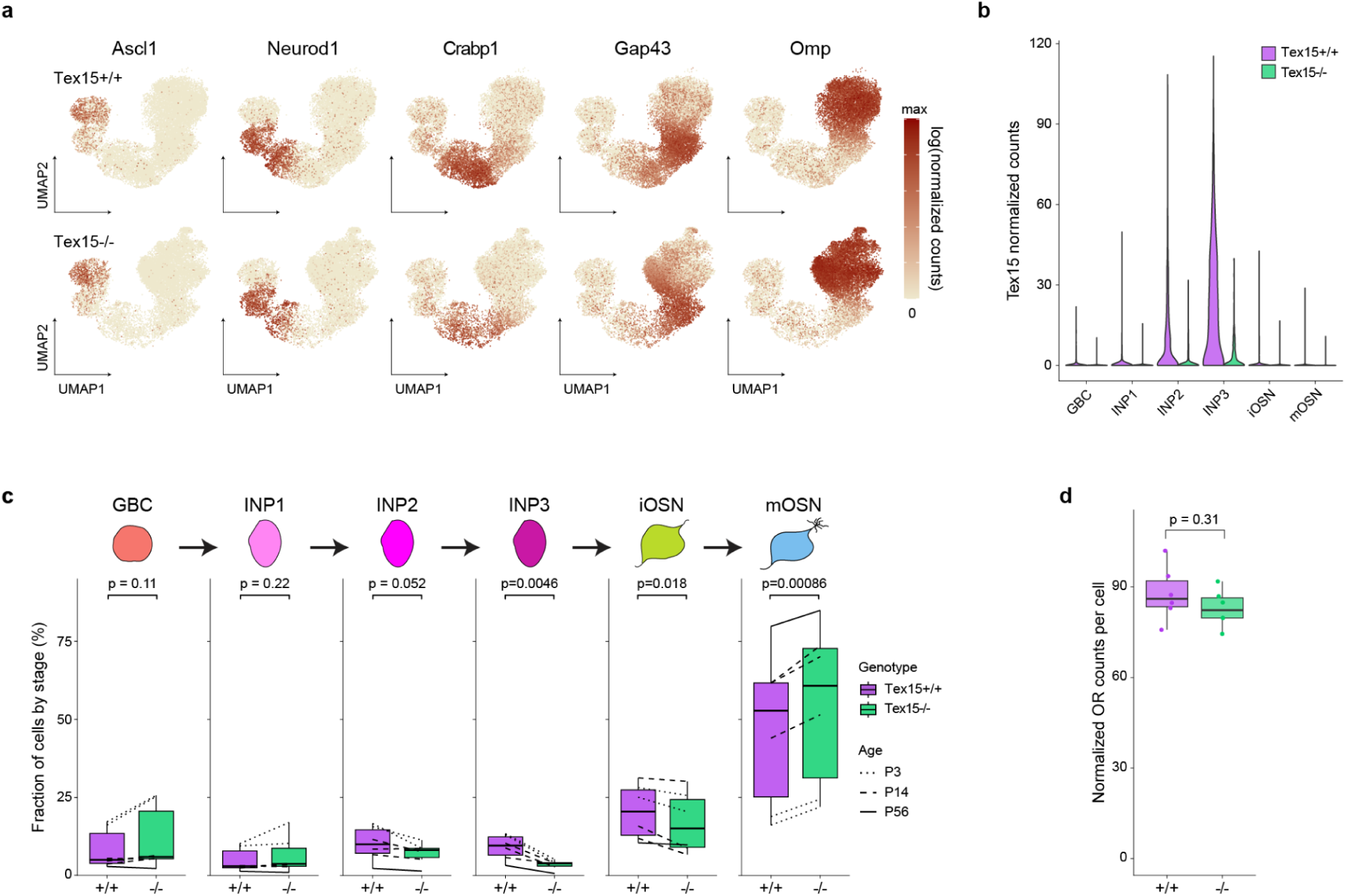
Single cell analysis of Tex15-/- and Tex15+/+ MOEs. **a**, UMAP projections in Tex15+/+ and Tex15-/- scRNAseq data showing log-normalized transcript levels of marker genes for mOSN differentiation. Ascl1 labels GBCs, Neurod1 labels INP1/2, Crabp1 labels INP2/3, Gap43 labels iOSNs, and OMP labels mOSNs. **b,** Normalized Tex15 counts per cell at each stage of differentiation. **c,** Abundance of cells at each stage of mOSN differentiation in Tex15+/+ and Tex15-/- samples. Lines connect paired samples and indicate the age of the mice. The Tex15-/- samples trend towards relatively few cells at the INP2 stage (p = 0.052), and then show a significant decrease at the INP3 (p = 4.57e-3) and iOSN (p = 1.79e-2) stages. However, they have a significant increase in the proportion in mOSNs (p = 8.58e-4) (paired t-test). **d,** Normalized OR counts in Tex15+/+ vs Tex15-/- mOSNs (t-test, p = 0.31).

**Extended Data 3:**
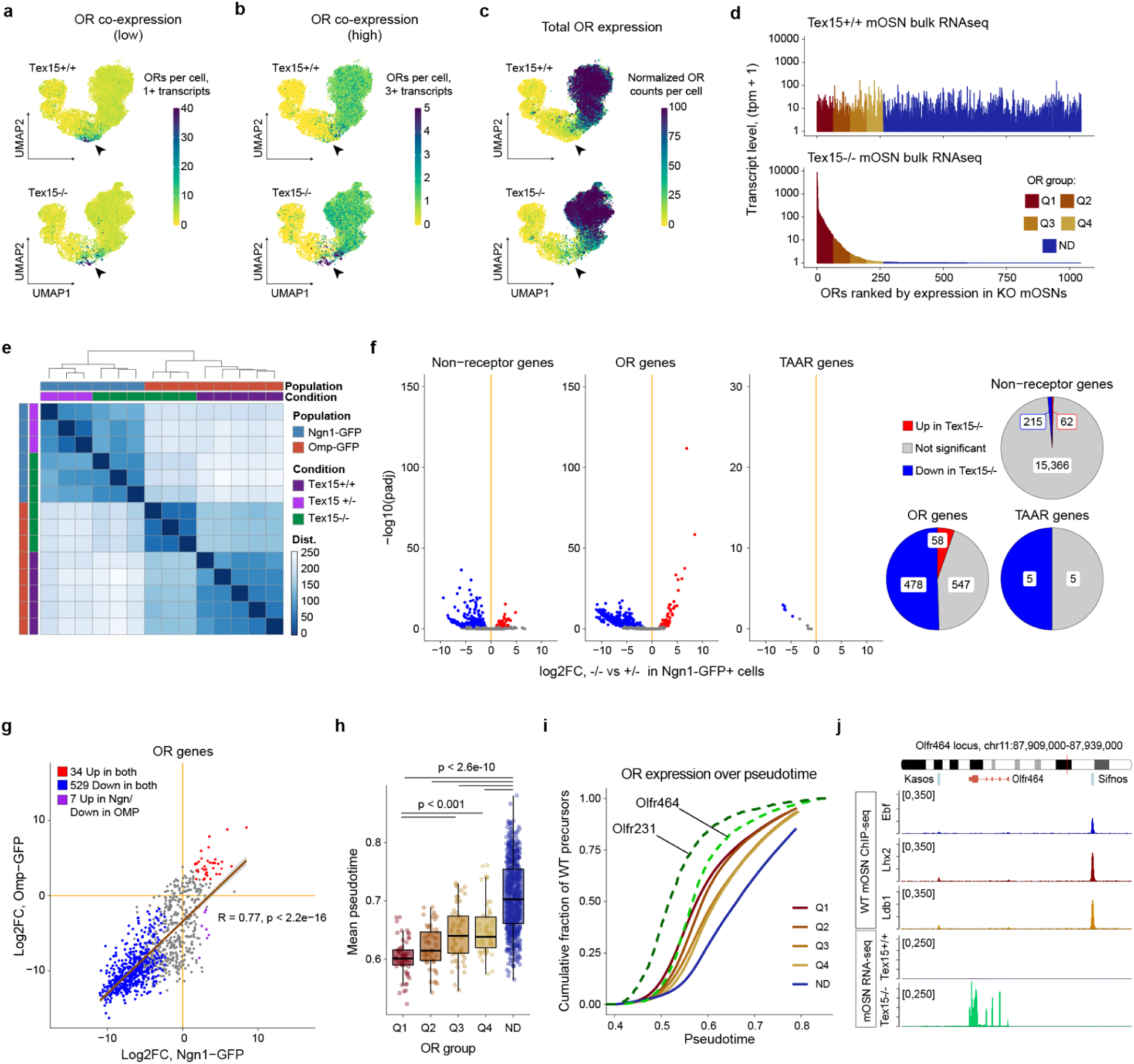
Early-expressed ORs dominate in Tex15-/- mice. **a**, Number of OR genes detected per cell using a low threshold for expression (1 or more transcripts per cell), reveals increased ORs co-expression in the Tex15-/- (at arrowhead) **b,** Number of OR genes detected per cell using a higher threshold for expression (3 or more transcripts per cell), reveals Tex15-/- INP3s co-expressing multiple ORs at unusually high levels. **c,** UMAP projection showing normalized OR counts per cell. **d,** Log-scaled OR transcripts levels (TPM + 1) sorted by level in Tex15-/- mOSNs. **e,** Heatmap showing euclidean distance between samples for top-1000 differentially expressed genes in Ngn-Gfp+ cells (INPs/iOSNs) and OMP-Gfp+ cells (mOSNs) **f,** Change in transcript levels between Tex15-/- and Tex15+/+ Ngn-Gfp+ cells. ORs and non-olfactory receptor genes are plotted separately. (right) summary of differentially expressed genes from each group (padj < 0.05 for a >50% change in expression, Wald test).**g,** Comparison of log2 fold change in OR gene transcript levels between Ngn1-GFP and OMP-GFP in Tex15-/- mice. **h,** The mean pseudotime value, corresponding to timing of OR expression, for each OR expression group. The Q1 ORs are expressed significantly earlier and the ND ORs are expressed significantly later than ORs in most other groups. **i,** The cumulative distribution of OR-expressing cells in wildtype precursors reveals that cells expressing Q1 ORs are abundant earlier in pseudotime than other groups, and the ND ORs appear relatively late. *Olfr231* expressing cells appear very early, even when compared to other Q1 ORs. In contrast, *Olfr464* resembles other Q1 ORs. **j,** *Olfr464* is located between two Greek Islands. ChIP-seq data from Monahan et al 2017 and Monahan et al 2019.

**Extended Data 4:**
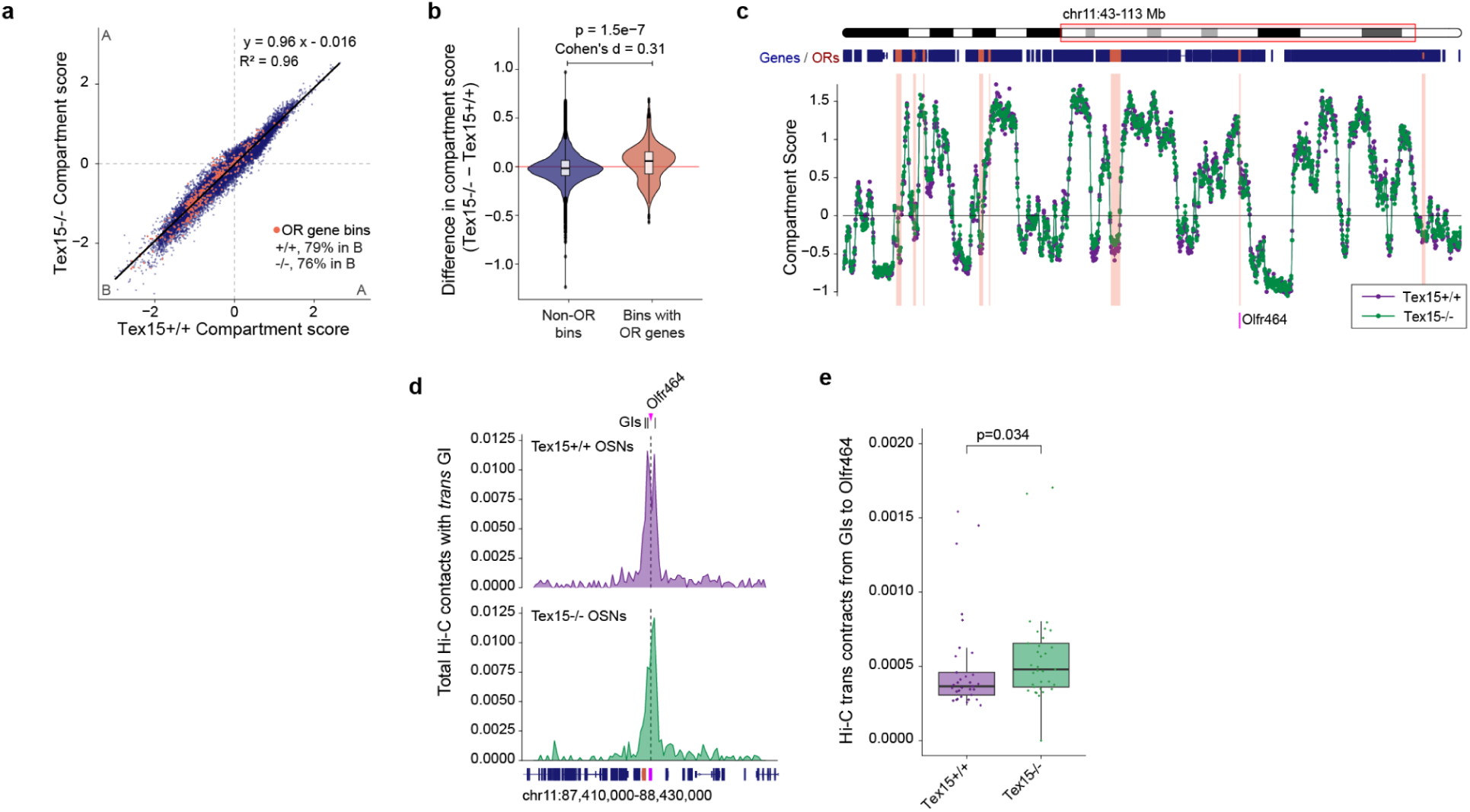
Genome organization and enhancer hubs in Tex15-/- mOSNs, **a**, Compartment scores for the whole genome (blue) and OR cluster loci (coral) are similar between Tex15+/+ and Tex15-/- mOSNS. 50 Kb bins. 79% of OR bins have a negative compartment score value (indicating compartment B) in the Tex15+/+ and 76% in the Tex15-/-. **b,** Bins that overlap OR genes exhibit a small but significant increase in compartment scores in Tex15-/- mOSNs (p=1.5e-7, t-test, Cohen’s d = 0.31). **c,** Signal tracks showing similar compartment scores across chromosome 11, including the region around *Olfr464*. **d,** The total Hi-C contacts made across the *Olfr464* locus by GIs located on a different chromosome (*trans* interchromosomal contacts) in Tex15+/+ and Tex15-/- mOSNs. **i,** Summed *trans* Hi-C contacts between GIs and the *Olfr464* locus in Tex15+/+ and Tex15-/- mOSNs. 10 Kb resolution. **e,** Hi-C contacts between *Olfr464* and each *trans* GI (Wilcoxon, p=0.034)

**Extended Data 5:**
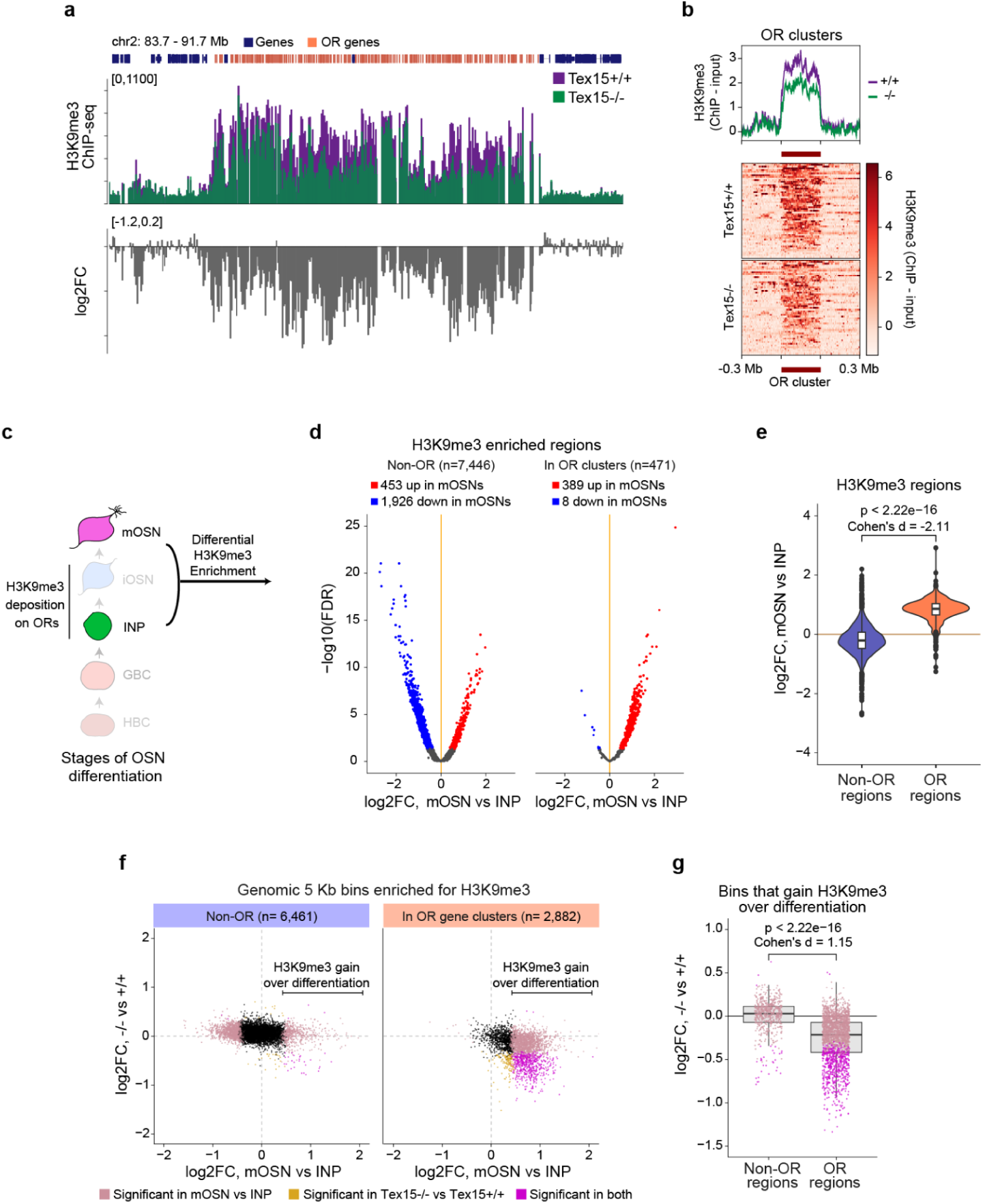
H3K9me3 marks on OR gene clusters. **a**, H3K9me3 ChIP-seq signal and log2 fold change over an OR gene cluster on chromosome 2. **b,** Heatmap showing H3K9me3 signal over OR gene clusters and flanking regions. Each row is an OR cluster, which are all scaled to be the same width. The plot above shows the average signal over these regions. **c,** Heatmap showing H3K9me3 signal over OR clusters genes and flanking regions. Each row is an OR gene cluster, which are all scaled to be the same width. The plot above shows the average signal over these regions. Only clusters over 100 Kb in size are included. **c,** Schematic illustrating the selection of cell stages to evaluate the gain of H3K9me3 on OR genes over differentiation. **d,** Differential enrichment analysis for H3K9me3 ChIP from mOSNs and INPs from Bashkirova et al 2023 (n=2 per stage). H3K9me3 enriched regions are split based upon overlap with OR gene clusters. Regions with a significant increase in H3K9me3 signal are in red, decrease in blue (FDR < 0.05). Nearly all regions that overlap an OR cluster gain H3K9me3 over differentiation. **e,** Fold change in ChIP signal for H3K9me3 regions grouped based on OR gene cluster overlap. OR regions show a significant gain in H3K9me3 signal over differentiation compared to non-OR regions (p < 2.22e-16, t-test, Cohen’s d = 2.25). **f,** Comparison of H3K9me3 fold change over differentiation versus fold change in Tex15-/-. All 5 Kb bins enriched for H3K9me3 in either condition are included. Bins are split based upon overlap with OR gene clusters. Bins with significant change in H3K9me3 are colored as indicated (FDR < 0.05). **g,** Fold change in H3K9me3 on 5 Kb bins that exhibit a significant increase in ChIP signal over differentiation. Bins overlapping OR gene clusters show a significant reduction in H3K9me3 signal compared to non-OR regions (p < 2.22e-16, Cohen’s d = 1.15). Panels **a,b** show data from a representative experiment.

**Extended Data Figure 6:**
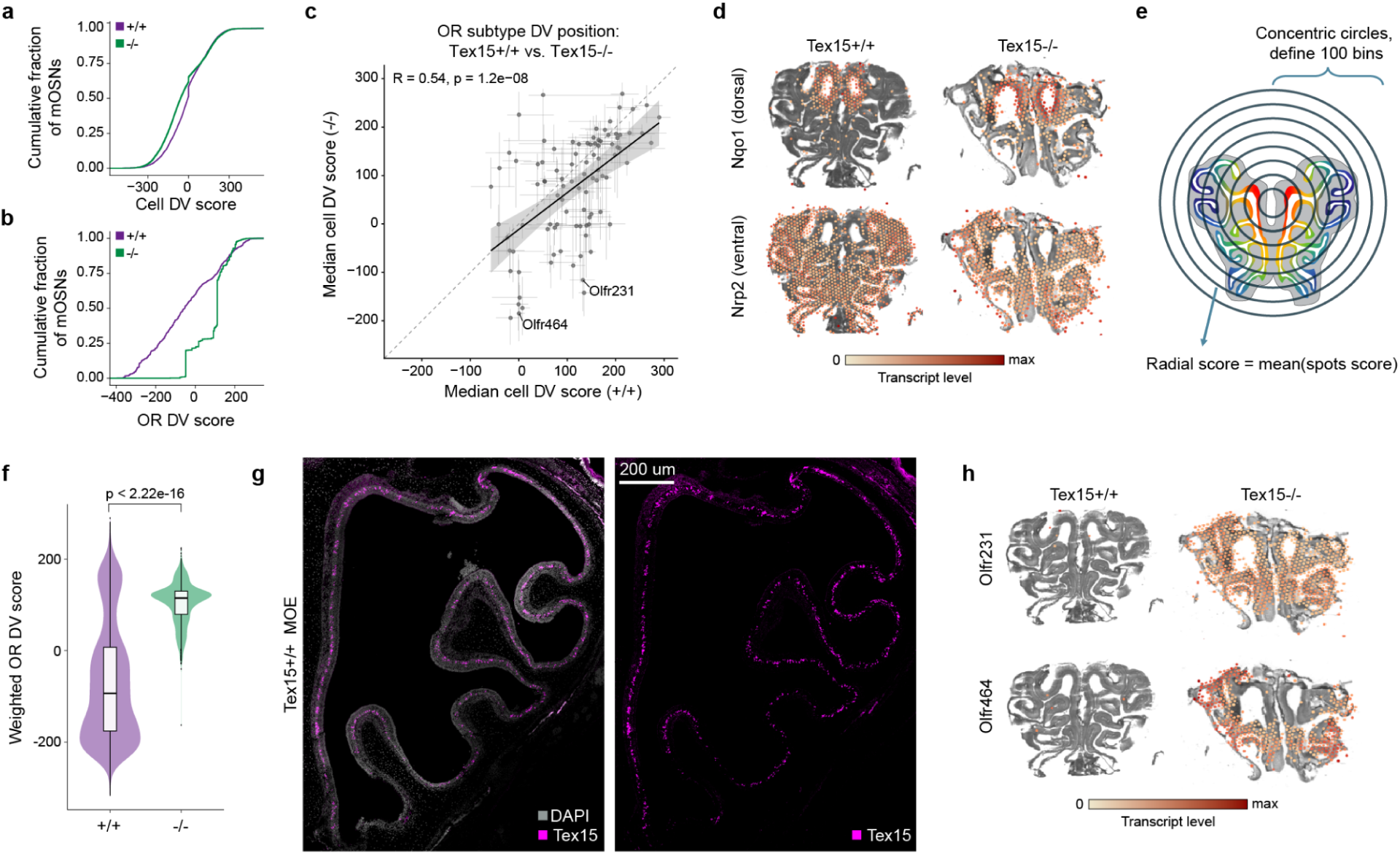
Spatial Pattern of MOE is disrupted in Tex15-/-. **a**, Cumulative fraction of Tex15+/+ and Tex15-/- mOSNs across the range of Cell DV scores. **b,** Cumulative fraction of Tex15+/+ and Tex15-/- mOSNs across the range of OR DV scores. **c,** Median cell DV score for mOSNs subtypes from Tex15+/+ and Tex15-/- mice. Only subtypes represented by at least 3 cells in the Tex15+/+ and Tex15-/- are included. Lines represent interquartile range. (R=0.54, p=1.2e-8, Pearson). **d,** Transcript levels of Nqo1, a dorsal marker gene, and Nrp2, a ventral marker gene, within each 55 um spot detected by spatial transcriptomics of a Tex15+/+ and Tex15-/- MOE representative section from LoupeBrowser. **e,** Schematic illustrating radial analysis strategy used to measure transcript levels along the DV axis across multiple slides. The “radial score” is calculated based upon the average of spot scores at the same radial distance from the center of the tissue, and then expressed as relative distance from center. **f,** The distribution of OR DV scores for spots on slides for Tex15+/+ (n=6) and Tex15-/- (n=3) MOEs. **g,** Tilescan showing Tex15 (magenta) immunoreactivity across half an MOE. Nuclei are labeled with DAPI (gray). Scale bar = 200 µm. **h,** *Olfr231* and *Olfr464* transcript levels within each 55 um spot of a Tex15+/+ and Tex15-/- MOE representative section from Loupe Browser.

## Author Contributions

N.Y. and K.M. designed the study and wrote the manuscript. K.M. conducted the bulk-RNA seq experiments and Hi-C experiments. N.Y. and J.K. conducted the single-cell experiments. N.Y. conducted the spatial transcriptomics experiments. N.Y. and A.I. conducted the ChIP-seq experiments. J.D. and N.B.G. conducted the RNAscope experiments. H.P. conducted the immunofluorescence experiments. D.B., S.R.D, K.M., and N.Y, designed and conducted single-cell analyses. C.S., J.Y. and N.Y., designed and conducted spatial transcriptomics analyses. P.J.W. provided the Tex15 null allele as well as technical assistance related to this allele. All other analyses were completed by K.M. and N.Y. with consultation and support from J.D. and D.B.

## Acknowledgements

We thank Paige Kramer for assistance with mouse colony management and help with preparing samples for fluorescent-activated cell sorting. We also thank Mackenzie Leung, Mallika Ravi, Pavithra Veera for assistance with mouse colony management. We acknowledge Arthur Roberts, Ankit Saxena, and the Rutgers Cancer Institute of New Jersey Flow Cytometry and Cell Sorting Shared Resource - supported, in part, with funding from NCI-CCSG P30CA072720-5921 - for flow cytometry and cell sorting services. We also thank Nadia Propp, Hemali Phatnani, and members of the Phatnani lab and the New York Genome Center for sharing their lab spaces and equipment for the spatial transcriptomics experiments. We acknowledge the School of Arts and Science-Human Genetics Institute of NJ (SAS-HGINJ) Imaging Core Facility and its director - Dr Zainab Tanvir - for microscopy services and support. We would also like to thank Eirene Markenscoff-Papadimitriou and Elizabeth Josie Clowney for their careful feedback on the manuscript. N.Y. was funded by the F31 pre-doctoral fellowship DC021641. This project was funded by startup funding from the School of Arts and Sciences at Rutgers, the State University of New Jersey and the Duncan and Nancy MacMillan Faculty Development fund, as well as R35GM146901 and the Rita Allen Scholars Fund (K.M). J.Y would like to acknowledge the startup funding from the School of Arts and Sciences and the Human Genetics Institute of New Jersey at Rutgers, The State University of New Jersey. P.J.W. would like to acknowledge R35GM153384, and P50HD068157. The content is solely the responsibility of the authors and does not necessarily represent the official views of the NIH.

## Data Availability

All newly generated data used in this study is publicly available in GEO accession number GSE341836. For gene expression across cell stages in Fig 1/Extended Data Figure 1, we used previously published data from Pourmorady et al 2024^14^, series GSE230380. For Shannon diversity index comparison in Figure 1i among published bulk RNA-seq data sets we used data available in GSE158730 from Bashkirova et al 2023, ^17^ for NfiABX knockout, GSE271029 from Dufflé et al 2025^38^ for Mex3a null, GSE325634 from Escamilla-Del-Arenal et al 2026^34^ for Cbx swap, and GSE277322 from Bao et al 2025^41^ for Trim66 null dataset. For comparison of Tex15 single knockout and Tex15 and Ldb1 double knockout bulk-RNA seq data to Ldb1 knockout bulk-RNA seq data in Figure 3, Extended data Figure 4, we used previously published Ldb1 knockout bulk-RNA seq data from Monahan et al 2019^25^, series GSE112153. FACS isolated mature OSN Hi-C data and Ldb1 ChIP-seq used in Figure 3 and Extended Figure 3j and Extended Data Figure 4 has been previously published in Monahan et al 2019 ^25^ and the data is publicly available in https://data.4dnucleome.org/ under accession number: 4DNESEPDL6KY. WT mOSN ChIP-seq data used in Extended Data Figure 3j is publicly available from Monahan et al 2017^42^ at GSE93570. For the pseudotime analysis for early transcription of ORs, we used data from Brann et al 2026^28^, publically available under GSE297068. For the heterochromatin deposition across cell stages data in Extended Data Figure 5, we used previously published data from Bashkirova et al 2023^17^, publicly available under GSE158730.

## Code availability

All code is publicly available in a GitHub repository (https://github.com/MonahanLab/Yusuf_2026), together with formatted reports from R workflow, additional data tables, and annotation files.

## Methods

Further information and requests for resources and reagents should be directed to and will be fulfilled by the Lead Contact, Kevin Monahan. No statistical methods were used to predetermine sample size. The experiments were not randomized and investigators were not blinded to allocation during experiments and outcome assessment.

### Mice

Mice were treated in compliance with the rules and regulations of Rutgers University - New Brunswick - IACUC under protocol number 2020008. All mice were housed in Specific Pathogen Free facilities in ventilated cages with a 12-h light/dark cycle, Mice had ad libitum access to standard rodent chow and water. Mice were group housed by sex. All experiments were performed on dissected whole main olfactory epithelium (MOE) or on freshly isolated, fluorescent activated cell-sorted primary cells collected from whole main olfactory epithelium.

The Tex15 knock-out mouse line was generated and published at Yang et al 2008^18^. Tex15 mice were crossed with Ngn-gfp^16^ to label immediate neuronal precursors and immature olfactory sensory neurons (OSNs). Tex15 mice were crossed with Omp-IRES-GFP mice^24^ to label mature olfactory sensory neurons (mOSNs). Conditional deletion of Ldb1 in mOSNs was achieved by crossing Ldb1 conditional allele mice(Ldb1-fl: Ldb1^tm2Lmgd^) with Cre-inducible tdTomato and Omp-ires-cre mice^43^. Double knockout of Ldb1-fl and Tex15-/- was achieved by crossing that to a Tex15+/- mouse.

For postnatal day 3, postnatal day 15, and adult single cell RNA-seq experiments, cells were sorted from male and female mice and sorted using the live cell FACs protocol below. For adult bulk RNA-seq, single-cell spatial transcriptomics, Hi-C, ChIP-seq experiments, cells were sorted from male and female mice ranging in age from 6 to 23 weeks. Biological replicate samples were collected and processed separately from different mice and as much as possible, littermate control and null mice were used for the experiments.

### Fluorescent Activated Cell Sorting (FAC sorting)

FACs experiments were conducted at Rutgers Cancer Institute of New Jersey Flow Cytometry and Cell Sorting Shared Resource. Mice under P10 were sacrificed directly with cervical dislocation or mice over P10 mice were sacrificed using CO2 followed by cervical dislocation.

The main olfactory epithelium (MOE) was dissected and transferred to ice-cold EBSS. The MOE was cut into small pieces with a razor blade, and then dissociated with papain (Worthington Biochemical, LK003163). Diced tissue was added to papain-EBSS, with at most 1-2 adult MOEs/mL or 2-3 young MOEs/mL, and incubated for 30-40 min at 37°C on a rocking platform.

After 30-40 min, tissue was triturated 30 times, the supernatant containing dissociated cells was transferred to a new tube, and the cells were pelleted (300 rcf, 5 min, room temperature).

Remaining papain was inhibited by resuspending the cell pellet with Ovomucoid protease inhibitor solution diluted 1:10 in EBSS, and the dissociated cells were pelleted (300 rcf, 5 min, room temperature).

For live cell sorts for native ChIP-seq experiments, dissociated cells were washed once with sort media (PBS with 2% Fetal Bovine Serum), and then resuspended in sort media supplemented with 100 U/mL DNase I, 4 mM MgCl2, and 500 ng/mL DAPI. These cells were passed through a 40 uM cell strainer, and then FAC sorted. Live cells were selected by gating out DAPI positive cells.

For live cell sorts for single-cell experiments, dissociated cells were washed once with wash media (1xPBS with 10% Fetal Bovine Serum, 5ug/ml Actinomycin, 10ug/ml Anisomycin, 10uM of Triptolide, 25U RNAse inhibitor), and then resuspended in sort media (1xPBS with 0.2% Fetal Bovine Serum, 5ug/ml Actinomycin, 10ug/ml Anisomycin, 10uM of Triptolide, 25U RNAse inhibitor, 100 U/mL DNase I, 4 mM MgCl2, and 500 ng/mL DAPI) and collected in collection media^6^. These cells were passed through a 40 uM cell strainer, and then FAC sorted. Live cells were selected by gating out DAPI positive cells.

For formaldehyde-fixed cell sorts, dissociated cells were resuspended in PBS + 1% methanol-free formaldehyde (Pierce). Cells were fixed at room temperature for 5 min, and then fixation was quenched by adding 1/10th volume of 1.25M glycine. Fixed cells were pelleted (500 rcf, 5 min, room temperature), washed once with sort media, resuspended in sort media, passed through a 40 uM cell strainer, and then FAC sorted.

### Native ChIP-seq

The MOE from 9 - 23 weeks old mice of Tex15+/+ and Tex15-/- were dissected and native chromatin was prepared as described before in Magklara et al 2011^16^. Briefly, nuclei were extracted and digested with micrococcal nuclease (MNase, Sigma), and then the digested chromatin was used in chromatin immunoprecipitation assays. The antibodies used were specific to H3 monomethyl lysine-9 (ab8896). All ChIP experiments were performed multiple times using different chromatin preparations.

### Bulk RNA-Seq

Live sorted cells were pelleted (15 min, 800 rcf, 4°C), the supernatant was aspirated until 250 uL of media remained, and then the cell pellet was resuspended in 750 uL Trizol LS (ThermoFisher). Total RNA was extracted by adding 200 uL chloroform, vortexing for 15 s, incubating at room temperature for 2 min, then centrifugation at 12,000 rcf for 15 min at 4°C. The aqueous phase was collected and RNA was precipitated with isopropyl alcohol with 10 ug/mL linear acrylamide (ThermoFisher) added as a carrier. The RNA pellet was washed twice with 75% ethanol, dried, then resuspended in nuclease free water. 1 ug of RNA was DNase treated using the TURBO DNA-free Kit (ThermoFisher) according to manufacturer’s instructions. RNA-seq libraries were prepared from DNase-treated RNA using a TruSeq Stranded Total RNA with Ribo-Zero Gold Set B kit (Illumina RS-1222302).

### Spatial Transcriptomics

Whole Tex15-/- and Tex15+/+ MOE of 6-7 weeks of age tissue was extracted, flash frozen with isopentane, and prepared for sectioning according to the fresh frozen tissue protocol outlined by 10x Cytassist Spatial Gene Expression for Fresh Frozen Tissue Preparation guide CG000636, Rev A. 10 μm cryosections of tissue were prepared in a clean, RNA-free cryostat on Superfrost slides. Sections were prepared for Hematoxylin & Eosin staining before starting Cytassist workflow to ensure quality and intactness of tissue using 10x Cytassist protocol number CG000614, Rev A. We prepared spatial transcriptome libraries from 6-7 weeks old adult wildtype and Tex15 knockout MOEs using Visium Cytassist Spatial Gene Expression workflow v4 (CG000495, Rev G). This Cytassist-based method has spatial spots that capture multiple cells within a 55 μm diameter and 45 μm distance between spots. Barcoded cDNA libraries of tissue sections were generated using the Spatial Gene Expression Reagent Kit (10x Genomics, PN-1000521). Data were demultiplexed and processed using SpaceRanger v4.0.1. Reads were aligned to the mm10 2020-A reference mouse transcriptome (10x Genomics) and using 10x mouse probe set 1 and OR transcriptome from Barnes et al 2020^2^.

### Single Cell Genomics

MOE was dissected at postnatal day 3, 14 or 56 from *Tex15-/-* and *Tex15+/+* littermate controls. Single cell RNA-seq libraries were generated using either Chromium single cell 3’ reagent kits v2 (adult replicates - PN-120267), v3.1 (both postnatal day 3 and 14 replicates PN-1000269, using 10x Chromium single cell 3’ protocol CG000315, Rev F) or v4.0 (postnatal day 3, PN-1000686, using 10x Chromium single cell 3’ protocol v4.0 CG000731, Rev B).

### Hi-C

In situ Hi-C libraries were prepared exactly as described in Monahan et al 2019^25^.

### Sequencing

Sequencing libraries were profiled on a Bioanalyzer 2100 using a high sensitivity DNA kit (Agilent) or Bioanalyzer Tapestation using a D5000 or D1000 DNA kit . Library concentration was determined by KAPA assay (KAPA Biosystems) or with Collibri™ Library Quantification Kit. Libraries were multiplexed and the pooled libraries were sequenced in either Illumina NovaSeq X or NextSeq 500.

### Immunofluorescence

MOE was dissected at postnatal day 7 and fixed in 4% PFA for 30 min on ice prior to being embedded in OCT. Coronal cryosections were taken at a thickness of 12 to 14 um and then air dried for 10 min. Slides were fixed with 4% PFA for 10 min. After fixation, slides were washed with PBST (PBS with 0.1% Triton X-100), blocked in PBST-DS (PBST +4% donkey serum). Slides were stained with primary antibody (Tex15, diluted 1:10, Gfp, diluted 1:1000, OMP, diluted 1:1000) in PBST-DS overnight at 4°C. Slides were then washed, stained with DAPI (2.5 ug/mL) and secondary antibody (Cy3 goat anti-mouse, diluted 1:1000, donkey a-rabbit conjugated to Alexa-488, diluted 1:1000, ThermoFisher) in PBST-DS for 1 hr, washed, and then mounted with Vectashield and imaged. Tex15 antibody was used from Boutros Ghali et al 2025^40^.

### Microscopy/Imaging

Confocal Images were collected with a Leica Confocal TCS SP8 built on a DMi8 microscope in the SAS-HGINJ Core Facility. Images were processed in the LAS X software associated with Leica microscope and Fiji software.

### AI Statement

No AI was used for generating any new concepts, experiments, text or code for this manuscript. This manuscript was proofread, refined to fit word count using Claude AI after the initial manuscript and ideas were written out by K.M. and N.Y and several rounds of comments were given by all authors of this manuscript. Claude AI also checked whether text and figure references were accurate, and both K.M. and N.Y. looked through all Claude AI suggestions before approving a small number of updates. After all data analysis was completed for this manuscript, the R-code shared on github was formatted for clarity using Claude AI.

### Statistical Test Statement

For each figure, the specific name of the test has been included in the legend or in the text. The code used for each statistical test has been included in our publicly available github repository.

## Analysis Methods

### Bulk RNA-Seq Analysis

Data processing and analysis was performed as previously described Monahan et al 2017^42^. Adaptor sequences were removed from raw sequencing data with CutAdapt version 5.2.

RNA-seq reads were aligned with STAR using a custom transcriptome annotation file based upon ENSEMBL release GRCm38.102, but updated to include expert curated odorant receptor gene annotation Barnes et al 2020 and Dietschi et al 2022^2,44^. SAMtools was used to select uniquely aligning reads by removing reads with alignment quality below 30 (-q30). RNA-seq data were analysed in R with the DESeq2 package, using a 1.5 fold change threshold for the p-value calculation, with rowmeans threshold set to 2, allowing for 1,088 of 1,141 protein-coding ORs to be detected. Differentially expressed genes were identified on the basis of an adjusted p-value of less than 0.05 for this fold change threshold. DESeq2 was also used to calculate normalized transcript levels as fragments per kilobase per million, which were then used to calculate transcripts per million for each gene. Analysis and plots including all genes for Figure 1a, Extended Figure 1b, Figure 2f uses previously published data from Pourmorady et al 2024^14^. All other bulk-RNA-seq analysis uses protein coding genes only for both olfactory receptor and non-olfactory receptor genes. Similar results are obtained if pseudogenes and other noncoding transcripts are included.

### Single Cell RNA-Seq Analysis

scRNA-seq fastq files were uniformed processed with a common Nextflow pipeline that ran Cell Ranger (v9.0.1) with a reference based on Ensembl v102 (GRCm10) genome and custom postprocessing to remove double-counted UMIs^6,28^. Data integration and identification of OSN lineage cells was performed using scvi-tools^45^. In brief, OR genes were excluded and the top 3,000 variable genes were identified using scVI’s poisson_gene_selection function. The raw counts for these genes were then used to train a scVI model (n_hidden: 128, n_latent: 30, n_layers: 2, dropout_rate: 0.2, gene_likelihood: ‘‘nb’’), with a batch key for each replicate, 10x chemistry as a categorical covariate, and total counts as a continuous covariate. A nearest neighbor graph (n_neighbors=25) of the resulting scVI latent embeddings was used for visualization via UMAP and clustered via the Leiden algorithm (resolution=1.6).

Clusters corresponding to OSN lineage cells were manually identified and higher quality non-doublet cells in these clusters with at least 500 genes and less than 20% of UMIs from mitochondrial genes were subclustered, and the above steps starting from variable gene identification were rerun. Similar custom methods were used for clustering in Brann et al 2026^28^. A nearest neighbor graph (n_neighbors=15) of the resulting latent space was constructed, clustered via the Leiden algorithm (resolution=0.6), and visualized via UMAP (min_dist=0.6).

The resulting clusters were manually annotated based on known marker genes (e.g. *Ascl1, Neurog1, Neurod1, Gng8, Lhx2, Hdac2, Gap43, Stmn2, Gnal, Adcy3, Omp*). Immature and mature OSNs were further iteratively subclustered, gene expression program usages were calculated by applying the GEP loadings identified in Tsukahara et al 2021^6^, and OR expression in each cell was evaluated based on the expression of functional OR genes at given thresholds (e.g. at least 1 or 3 UMIs).

### Hi-C Analysis

Fastq files were aligned to the mouse genome (mm10) using the distiller-nf pipeline version 0.3.4 (https://github.com/open2c/distiller-nf). Uniquely mapped reads (mapq > 30) were retained, and duplicate reads were discarded. Cooler was then used to bin contacts into matrices and calculate balanced contact values using default parameters except setting “--mad-max 7” and “--min-nnz 5”, which improves coverage of OR cluster regions. Data pooled from two biological replicates were analysed together, after the results of analyses of individual replicates had been confirmed to be similar. Balanced Hi-C contacts were analyzed and visualized in R using Genova v.1.0^46^.

### Native ChIP-Seq Analysis

Sequencing reads were pre-processed by trimming adapters with Cutadapt version 5.2^47^. Trimmed reads were then aligned to the mouse genome (mm10) using Bowtie2 version 2.3.2^48^ with default settings except for maximum insert size set to 1000 (-X 1000), which allows larger fragments to be mapped. Duplicate reads were removed with Samtools version 1.19.2^49^, and then the uniquely mapped, properly paired reads were selected with Samtools (-q 30 -f 3).

Native ChIP heatmaps were generated with deeptools^50^ with OR gene bodies re-scaled to 6kb and showing 2kb flanking on each side.

Genome wide analysis of H3K9me3 ChIPseq data was performed in R with csaw^51^. Regions enriched for H3K9me3 were identified by performing a binned analysis, in which reads were mapped to 5 Kb and 50 Kb bins spanning the genome. The 50 Kb bins were used to calculate a global background, then enriched 5 Kb bins were identified by filtering for greater than 3-fold enriched relative to the global background. The global background for each library was used to generate a normalization factor to compare read counts between libraries. EdgeR version 4.10.1^52^ was then used to identify bins that were differentially enriched between conditions. The raw p-value for each bin was then corrected by calculating and false discovery rates (FDR). Since H3K9me3 marks tend to cover broad areas, groups of nearby bins were joined into H3K9me3-enriched “regions”, and csaw was used to calculate a region-based p-value and FDR.

To compare the fold change in H3K9me3 signal between cell-stage and *Tex15-/-* experiments, each experiment was first analyzed separately, including the identification of enriched 5 Kb bins and the calculation of fold change in H3K9me3 between conditions. Then the 5 Kb bins that were significantly enriched in both experiments were selected, and these were used to compare fold changes across experiments.

### Visium data processing and spot filtering

10x Genomics Visium spatial transcriptomics data from adult mouse main olfactory epithelium (MOE) sections were analysed in R (v4.3.1) using Seurat 5.5.1^53^ and Harmony 2.0.5^54^. The final dataset contained six wild-type and three Tex15 knockout MOE sections. Odorant receptor (OR) genes were defined using the OR annotation from Barnes et al 2020^2^. For each spot, the number of detected OR genes and total OR transcript counts were calculated from the Spatial assay counts. Spots were retained if they expressed at least two OR genes and at least three total OR transcripts. Dorsal and ventral non-OR marker genes were taken from Tuskahara et al 2021^6^. Marker genes were split by annotated direction, yielding 611 dorsal and 739 ventral markers before intersection with genes detected in the Seurat object. To compare dorsal-ventral patterning using radial profiling across MOE sections, spot coordinates were transformed into a normalized radial coordinate. For each section, the centroid of Zone 1 spots was used as the tissue center. Euclidean distance from this centroid was calculated for every spot and min-max normalized within each section to produce a radius from 0 to 1. Spots were assigned to 100 equally spaced radial bins. Mean scaled cell DV score and mean OR-weighted DV score were calculated within each bin for each section. Genotype-level radial profiles were generated by averaging the binned section-level profiles within wild-type and Tex15 knockout groups; shaded intervals show the standard error of the mean across sections.

